# Cerebral Small Vessel Disease genetic determinant TRIM47 controls brain homeostasis *via* the NRF2 antioxidant system

**DOI:** 10.1101/2024.10.08.616723

**Authors:** Cécile Duplàa, Valentin Delobel, Romain Boulestreau, Sébastien Rubin, Juliette Vaurs, Béatrice Jaspard-Vinassa, Muriel Busson, Cloé Combrouze, Carole Proust, Jean- Luc Morel, Bruno Bontempi, Aniket Mishra, Stéphanie Debette, Thierry Couffinhal, Claire Peghaire

**Affiliations:** Univ. Bordeaux, INSERM, Biologie des maladies cardiovasculaires, U1034, F-33600 Pessac, France; CNRS, IMN, UMR 5293, University Bordeaux, F-33000, Bordeaux, France; Univ. Bordeaux, INSERM, Bordeaux Population Health, U1219, F-33000 Bordeaux, France

## Abstract

Cerebral small vessel disease (cSVD) is a leading cause of stroke, cognitive decline and dementia, for which no specific mechanism-based treatments are available to date. Genome-wide and whole-exome association studies previously identified robust associations of common variants at chr17q25 with cSVD features on magnetic resonance imaging, with converging bioinformatic and experimental data for a causal involvement of *TRIM47*. Preliminary functional evaluation of TRIM47, an ubiquitin ligase enriched in brain endothelial cells (ECs), suggested its potential role in blood brain barrier (BBB) integrity. Here, we show that TRIM47 regulates brain EC resilience and adaptive responses to oxidative stress by binding to KEAP1, stabilizing NRF2 protein levels and promoting the NRF2 antioxidant signaling pathway. *In vivo*, *Trim47*-deficient mice exhibit downregulation of NRF2 target genes, BBB dysfunction, astrogliosis and cognitive impairments. Endothelial-specific deletion of *Trim47* recapitulates these phenotypes. Treatment with the NRF2 activator tert-butylhydroquinone normalized BBB integrity and cognitive function in *Trim47*-deficient mice, highlighting the role of endothelial TRIM47 in driving brain homeostasis through NRF2 pathway activation. This work indicates that loss of the protective TRIM47/NRF2 axis may increase the susceptibility to developing human cSVD and that targeting the TRIM47/NRF2 axis could be a promising therapeutic approach for vascular cognitive impairment and dementia.

## Introduction

Cerebral small vessel disease (cSVD) encompasses a range of pathological processes that damage arterioles, capillaries or venules supplying brain tissue. It is a leading cause of stroke (both ischemic and hemorrhagic) and the major pathological substrate underlying the vascular contribution to cognitive decline and dementia, including Alzheimer type dementia^1–4^. cSVD is highly prevalent in the general population with increasing age and is often covert, i.e. detectable on brain magnetic resonance imaging (MRI) before clinical symptoms manifest^5,6^, thus representing a major target for prevention^7^. Key MRI-features of cSVD include white matter hyperintensities (WMH), microbleeds, lacunes and perivascular spaces^8^, reflecting consequences of cSVD on the brain parenchyma.

Increasing age and high blood pressure are the strongest known risk factors for cSVD. However, while lowering blood pressure, and more broadly optimal management of vascular risk factors, is an important approach to prevent and slow down the progression of cSVD, it is insufficient^9,10^, with vascular risk factors explaining only a small proportion of WMH variability in older age^11^. Specific treatments for cSVD are currently lacking, partly because the molecular pathways driving its etiology remain poorly understood. Thus, understanding the precise molecular and cellular mechanisms underlying cSVD and its consequences on cognitive performance is an absolute prerequisite for identifying novel therapeutic targets.

Genomics is powerful tool to identify disease pathways and was repeatedly shown to provide a strong foundation for mechanistic studies and target discovery^12,13^. It has been estimated that genetic support for new drug efficacy more than doubles success rates in clinical trials^13–16^. Large collaborative genome-wide and whole-exome association studies on population-based cohort studies have identified numerous genetic risk variants associated with MRI-markers of cSVD^3,17–20^. The most significant association with WMH volume and a composite extreme-cSVD phenotype was described at chr17q25^3,21^. Within this gene-rich locus, summary-based Mendelian randomization and profiling of human loss-of-function allele carriers in UK Biobank pointed to *TRIM47* as the most plausible causal gene with evidence for loss-of-function mechanisms driving the association with cSVD severity^21^. Moreover, transcriptomic data showed an enrichment of *TRIM47* expression in brain blood vessels and endothelial cells (ECs) (https://markfsabbagh.shinyapps.io/vectrdb) and preliminary functional evaluation with siRNA targeting *TRIM47* showed increased endothelial permeability *in vitro*^21^. In the present study, we devised a multilayered experimental plan to decipher the biological mechanisms underlying the association of TRIM47 with cSVD.

TRIM47 belongs to a subfamily of E3 ubiquitin ligases involved in many cellular and physiological processes including cell proliferation, apoptosis, innate immunity and autophagy. The subfamily of E3 ubiquitin ligases regulates protein degradation, cell trafficking and is also involved in DNA repair^22,23^. TRIM47 was first identified as being overexpressed in astrocytoma^24^ and its role in cancers has been studied extensively. As an oncogene, TRIM47 promotes tumorigenesis, and is overexpressed in multiple cancers (breast^25^, ovarian^26^, colorectal^27^, glioma^28,29^, renal^30^, gastric cancers^31^). It is also used as a prognostic biomarker for several cancers^31–34^.

Previous work also demonstrated that TRIM47 regulates cellular functions associated with angiogenesis *in vitro*^28^, with its expression being upregulated under various pathological conditions, including inflammation, hypoxia, and oxidative stress in human umbilical vein endothelial cells (HUVEC)^35^. Recently, two studies provided additional evidence for a possible role of TRIM47 in vascular biology and more specifically its deleterious effect in inflammatory contexts. *In vivo*, deletion of *Trim47* was shown to alleviate inflammation in a mouse model of LPS-induced sepsis^35^, while overexpression of *Trim47* markedly accelerated cerebral ischemic injury though promoting apoptosis and inflammation in rats^36^.

In the present study, we demonstrate a major contribution of TRIM47 to brain endothelial cell homeostasis, blood brain barrier integrity, and cognitive function, through regulation of the antioxidant NRF2 pathway. Our data highlight the therapeutic potential of targeting the TRIM47/NRF2 protective axis in patients with cSVD and vascular cognitive impairment and dementia (VCID).

## Results

### TRIM47 is an important activator of the antioxidant NRF2 pathway in human brain endothelial cells

Our group, along with others, has recently reported that TRIM47 is not only expressed by various types of ECs, but is also particularly enriched in brain vessels and plays a crucial role in regulating ECs functions *in vitro*^21,28,35^. To identify the specific pathways regulated by TRIM47 in brain ECs, we performed bulk RNA-sequencing on human brain microvascular endothelial cells (HBMEC) treated with either control siRNA (siCtl) or *TRIM47*- targeting siRNA (si*TRIM47*) for 72 h. Transcriptomic data analysis revealed that 208 genes were downregulated while 214 genes were upregulated following *TRIM47* depletion (Fig. 1a). Pathway enrichment analysis revealed that the genes regulated by TRIM47, particularly those downregulated in *TRIM47*-deficient cells, were associated with the nuclear erythroid 2-related factor 2 (NRF2) pathway, oxidative stress and terms related to NRF2 (Nuclear receptors metapathway, NRF2ARE regulation, NRF2 transcriptional activation, NRF2 survival signaling) (Fig. 1b). Interestingly, *TRIM47* depletion did not affect the mRNA levels of the transcription factor *NFE2L2* (the gene encoding NRF2) and Kelch-like ECH-associated protein 1 (*KEAP1*), a regulator of NRF2, but it significantly downregulated key NRF2 target genes including *HMOX1*, *NQO1* and *GCLM* (Fig. 1c). Further time course experiments showed that inhibition of TRIM47 expression in HBMEC by siRNA treatment reduced *NQO1* and *HMOX1* (HO1) mRNA levels over 24, 48 and 72 h (Supplementary Fig. 1a-c) with a corresponding decrease in HO1 protein levels (Fig. 1d).

**Fig. 1.**
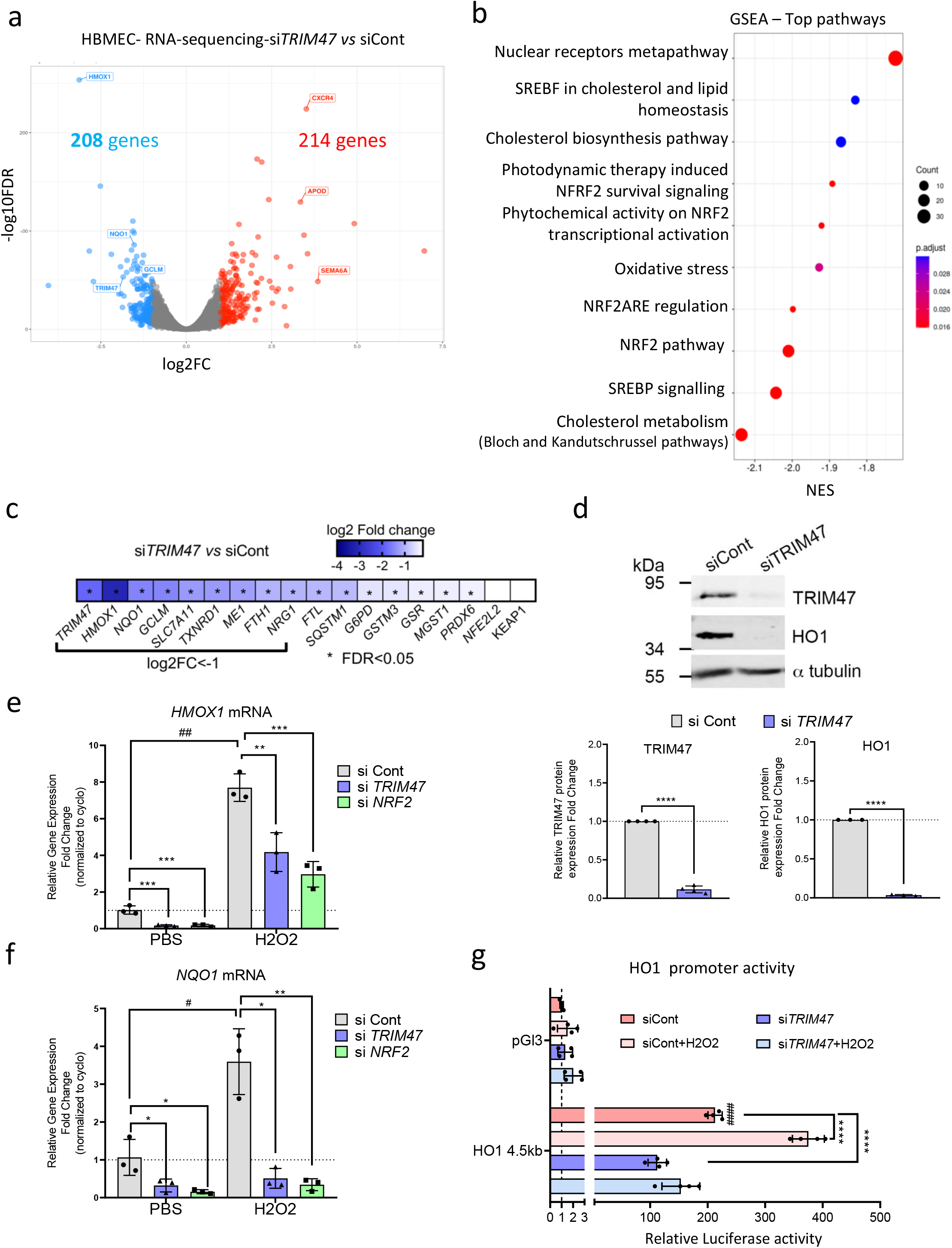
TRIM47 is an activator of the antioxidant NRF2 pathway in human brain endothelial cells **a.** RNA-sequencing was performed on HBMEC treated with control or TRIM47 siRNA for 72h. Volcano plot showing log2 fold change (FC) *vs* -log10 FDR of differentially expressed genes (DEG) in response to TRIM47 knockdown. Significant downregulated (log2FC<-1, in blue) and upregulated (log2FC>1, in red) genes were identified by edgeR analysis (FDR<0.05 and CPM >=5). **b.** Pathway enrichment analysis of DEG. Gene set enrichment analysis (Wikipathway) revealed a significant (p-adj=0.025) downregulation of genes associated with the NRF2 signalling pathway. Dotplot depicted the normalized enrichment scores (NES) for the top 10 pathways. The dots are colored by the adjusted p- value and their size is proportional with the size of the gene-set. **c.** Heatmap showing the expression of selected genes related to the NRF2 pathway in HBMEC treated with TRIM47 siRNA *vs* control. Data are expressed as log2FC. *FDR<0.05. **d.** Immunoblot and quantification of WB for TRIM47 and HO1 expression in control and TRIM47-deficient HBMEC treated with DMSO or MG132 for 5h. Data were normalized to α-tubulin (n=3 or 4 experiments). **** P<0.0001, Student’s t*-*test. **e-f.** qPCR analysis of (**e**) *HMOX1* and (**f**) *NQO1* gene expression in control, TRIM47 or NRF2 siRNA-treated HBMEC for 72h and treated with either PBS or H2O2 (200 uM for 3h). Data were normalized to *cyclophilin* (n=3 experiments). * P<0.05; ** P<0.01; *** P<0.001, One-way ANOVA, # P<0.05; ## P<0.01, Student’s t- test. **G.** HO1 promoter luciferase reporter assay. HeLa were co-transfected with HO1 promoter- luciferase construct (HO1-4.5bp) or a pGL3 empty vector with control or siRNA targeting TRIM47. Cells were treated with either PBS or H2O2 (200 uM for 1h). Luciferase activity was measured in cell lysates. Values represent the fold change in relative luciferase activity over the HO1-4.5kb + siControl condition and normalized to Renilla (n=4 experiments). **** P<0.0001 One-way ANOVA; #### P<0.0001 compared to pGL3 + siControl condition, Student’s t-test. All graphical data are mean ± s.d.

Additionally, *TRIM47* expression was upregulated in HBMEC but *NFE2L2* mRNA levels were not affected (Supplementary Fig. 1d-e) under oxidative stress conditions induced by H_2_O_2_, corroborating previous observations in HUVEC^35^. Crucially, TRIM47 was required to fully induce *HMOX1* and *NQO1* mRNA levels in both basal and oxidative stress conditions in HBMEC, mirroring the effects of *NRF2* siRNA (Fig. 1e-f). In line with these data, H_2_O_2_ treatment activated the HO1 promoter (HO1-4.5kb); *TRIM47* depletion caused a significant decrease in HO1 promoter activity in both baseline and oxidative stress conditions (Fig. 1g). Together, these data showed that TRIM47 is a key and novel activator of the NRF2 antioxidant pathway in human brain ECs.

### TRIM47 regulates NRF2 stability *via* its interaction with KEAP1 in brain endothelial cells

To uncover the precise mechanism by which TRIM47 regulates the NRF2 pathway, we investigated the potential interaction between TRIM47 and the transcription factor NRF2 in brain ECs using *in vitro* models. *TRIM47* overexpression (Supplementary Fig. 2a) led to a modest increase in *NFE2L2* expression (Supplementary Fig. 2b) and to a significant increase in the expression of NRF2 target genes *NQO1* and *HMOX1* (Fig. 2a-b). This induction was abolished by *NRF2* siRNA treatment showing that TRIM47’s effect on these genes is dependent on NRF2 (Fig. 2a-b).

**Fig. 2.**
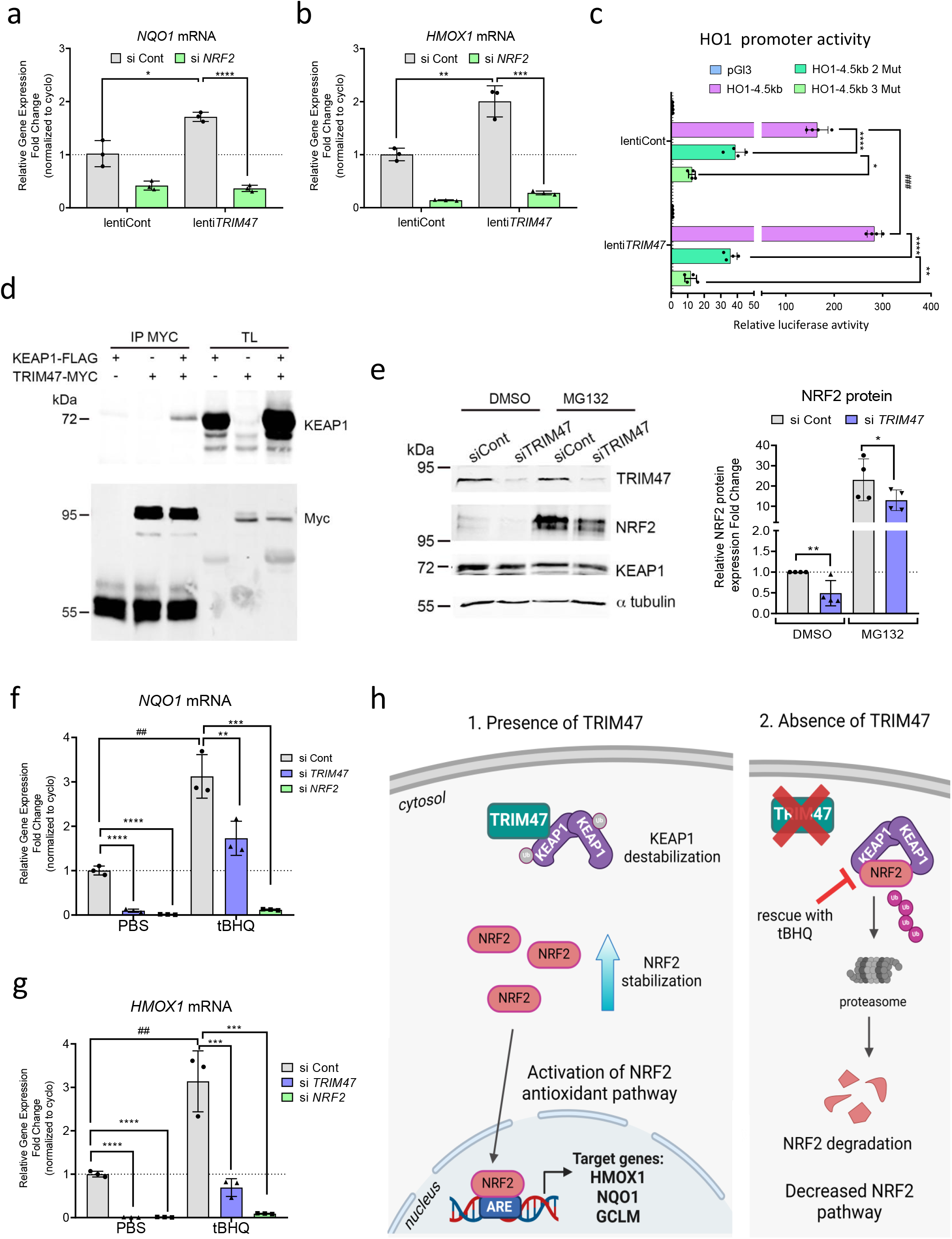
TRIM47 cooperates with NRF2 in driving antioxidant target genes. **a-b.** qPCR analysis of (**a**) *NQO1* and (**b**) *HMOX1* gene expression in siCont or siNRF2 treated HBMEC and co-transduced with a control or TRIM47 lentivirus for 24h. Data were normalized to *cyclophilin* (n=3 experiments). * P<0.05; ** P<0.01; *** P<0.001; **** P<0.0001, One-way ANOVA. **c.** HO1 promoter luciferase reporter assay. TRIM47 cDNA expression plasmid (TRIM47) or empty expression plasmid (pcDNA) were co-transfected with HO1 promoter-luciferase constructs (HO1-4.5bp, HO1- 4.5bp with 2 or 3 mutations) or a pGL3 empty vector in HeLa, and luciferase activity was measured. Values represent the fold change in relative luciferase activity over pGL3 vector (n=4 experiments). * P<0.05; ** P<0.01; **** P<0.0001, One-way ANOVA; ### P<0.001: compared to pGL3 + Lenti control condition, Student’s t-test. **d.** TRIM47 and KEAP1 interaction was assessed by Co-IP assay in whole cell lysates from HEK293. Cells were transfected with TRIM47-myc pcDNA, KEAP1-Flag or both expression plasmids. Lysates were immunoprecipitated with myc antibody and immuno-blotted for myc (TRIM47) and KEAP1. **e.** Immunoblot and quantification of WB for NRF2 and KEAP1 in control and TRIM47-deficient HBMEC treated with DMSO or MG132 for 5h (n=4 experiments). Data were normalized to α-tubulin.* P<0.05; ** P<0.01, Student’s t-test. **f-g.** qPCR analysis of (**f**) *NQO1* and (**g**) *HMOX1* gene expression in HBMEC treated with control, TRIM47 or NRF2 siRNA for 72h and with tBHQ (10µM) for 24h (n=3 experiments). ** P<0.01; *** P<0.001; **** P<0.0001, One-way ANOVA; ##, P<0.01 Student’s t*-*test. All graphical data are mean ± s.d. **h.** Model of TRIM47 activation of the antioxidant NRF2 pathway in endothelial cells. 1. In basal conditions, TRIM47 binds to KEAP1 in the cytoplasm and thus alleviates KEAP1-dependent NRF2 degradation and increases NRF2 protein stabilization that can translocate to the nucleus to drive the expression of NRF2-target genes (*HMOX1*, *NQO1*) promoting cellular antioxidant protection. 2. In the absence of TRIM47 (TRIM47 siRNA condition), KEAP1 strongly binds to NRF2 leading to its ubiquitination and degradation by proteosomal complex, thus leading to the decreased of the protective NRF2 pathway.

Given that NRF2 is the master regulator of antioxidant defense gene expression and directly drives the expression of HO1 (the best described NRF2 target and a direct NRF2 binding partner) and NQO1 by binding to antioxidant response elements (ARE), on their promoters (reviewed in^37^), we sought to determine whether TRIM47 requires these NRF2 DNA binding sites to increase HO1 expression. A luciferase assay in HeLa cells revealed that *TRIM47* overexpression activates the HO1 promoter (Fig. 2c). Mutations in the two ARE sites on the HO1 promoter (HO1-4.5kb 2 Mut, double mutant construct)^38,39^ blocked HO1 promoter transactivation, underscoring the necessity of NRF2 binding to ARE sites for TRIM47-mediated HO1 induction (Fig. 2c). Similar effects were also observed with a triple mutant construct with additional mutation in the E box of the HO1 promoter (HO1-4.5kb 3 Mut) which fully blocked HO1 promoter transactivation induced by TRIM47 (Fig. 2c).

The activity of NRF2, both under basal conditions and in response to stress, is tightly controlled. NRF2 protein levels are finely regulated by KEAP1, a master negative regulator that binds to NRF2, leading to its ubiquitination and subsequent degradation by the proteasome (reviewed in^37^). To investigate whether TRIM47 modulates the NRF2 pathway through the formation of a regulatory complex, we performed co-immunoprecipitation (Co-IP) assays. Our data revealed that TRIM47 interacts with KEAP1 in HeLa cells overexpressing both TRIM47-myc and KEAP1-Flag (Fig. 2d), but not with NRF2 itself (data not shown). Importantly, although TRIM47 depletion in HBMEC did not affect KEAP1 protein levels, Western blot analysis, with and without a proteasome inhibitor treatment (MG132), showed a significant decrease in NRF2 protein (Fig. 2e). This suggests that TRIM47 may prevent NRF2 protein degradation by binding to KEAP1. ECs were then treated with an NRF2 pathway activator, namely the tert-butylhydroquinone (tBHQ), which promotes NRF2 protein stability by destabilizing the KEAP1-NRF2 complex. This NRF2 pathway activator did not affect the mRNA levels *of TRIM47* (Supplementary Fig. 2d) and *NFE2L2* (Supplementary Fig. 2e). However, importantly, tBHQ was able to partly rescue the effect of *TRIM47* siRNA on *NQO1* (Fig. 2f) and *HMOX1* (Fig. 2g) expression, indicating that TRIM47 acts upstream of NRF2. As expected, tBHQ could not rescue the effects of *NRF2* siRNA on these target genes (Fig. 2f and g). These findings highlight TRIM47 as a key modulator of NRF2 stability through its interaction with KEAP1.

In summary, these data indicate that under basal conditions when TRIM47 is expressed in ECs, it binds to KEAP1 in the cytoplasm, thereby reducing NRF2 protein degradation and enhancing its stability. NRF2 can then accumulate in the cytoplasm and translocate to the nucleus and binds to ARE sites to induce the transcription of its antioxidant target genes. Conversely, when TRIM47 is repressed, KEAP1 is free to bind to NRF2, leading to its ubiquitination and degradation by the proteasome, which decreases the NRF2-mediated antioxidant pathway in ECs. The NRF2 pathway can be reactivated by the use of the NRF2 activator tBHQ as depicted in the model (Fig. 2h).

### TRIM47 protects from oxidative stress both *in vitro* and *in vivo*

Since TRIM47 promotes the antioxidant NRF2 pathway, we hypothesized that loss of TRIM47 in brain ECs might lead to a higher susceptibility to stress. Indeed, depletion of *TRIM47* in HBMEC led to a significant increase in oxidative stress as detected by a CellRox green dye. Additionally, *TRIM47* siRNA worsened the effect of two oxidative stress inducers, tert-butyl hydroperoxide (TBHP) and Angiotensin II (Supplementary Fig. 3a and b). Notably, activation of the NRF2 pathway by stabilizing NRF2 protein with the tBHQ compound was able to dampen the oxidative stress induced by *TRIM47* depletion under both basal and stress conditions (Fig. 3a-b). This indicates that TRIM47 contributes to antioxidant defenses in HBMEC, at least in part by activating the NRF2 pathway.

**Fig. 3.**
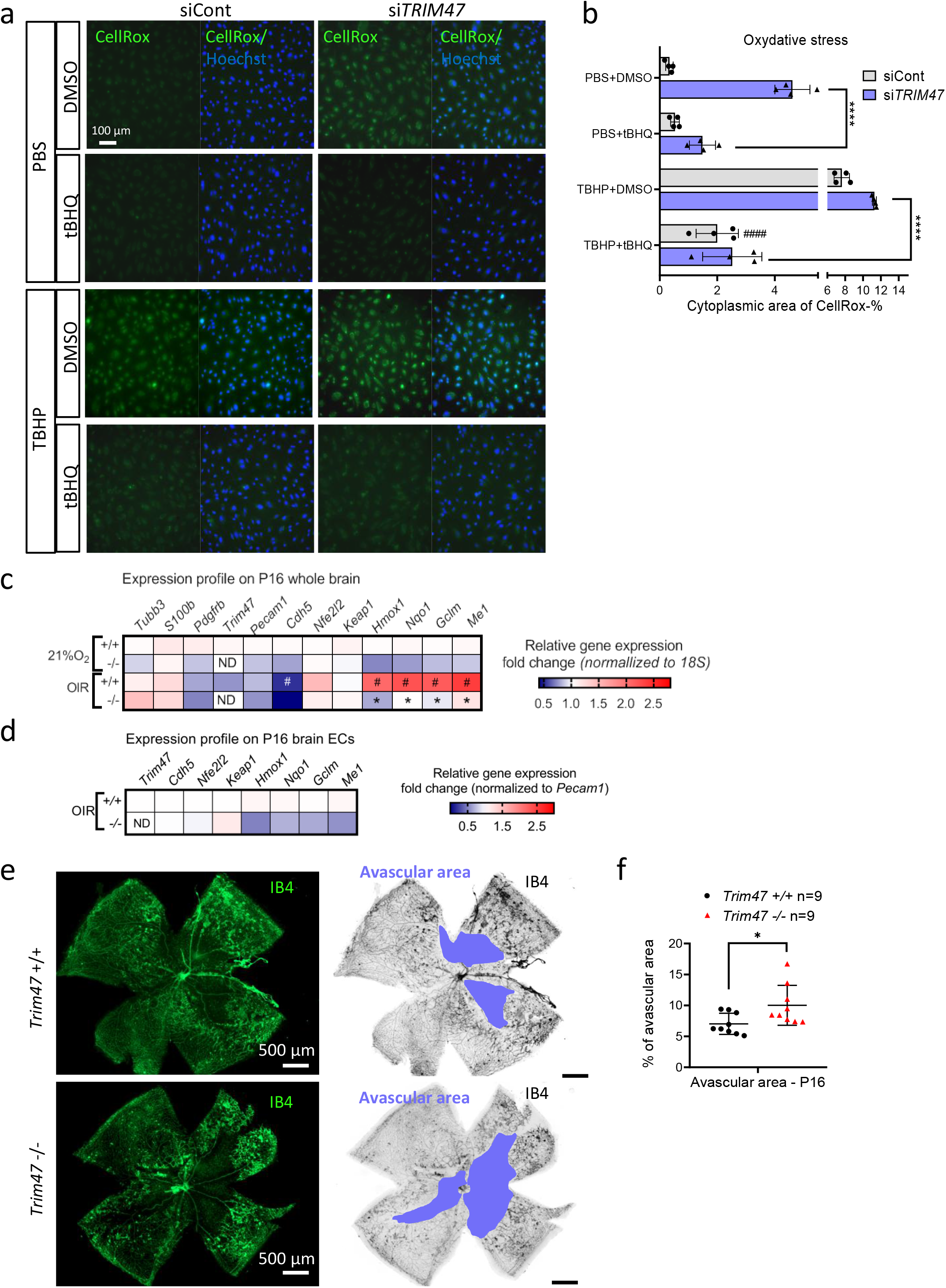
TRIM47 protects from oxidative stress *in vitro* and *in vivo* **a-b.** Representative immunofluorescence image (**a**) and quantification (**b**) of CellRox dye (green, marker of oxidative stress) in HBMEC transfected with siCont or si*TRIM47* for 24 h, then pre-treated with DMSO or tBHQ (10 µM for 24 h) and treated with PBS or tert-butyl hydroperoxide (TBHP, 200 µM for 1 h); nuclei are identified by Hoechst (blue). Scale bar 100 µm. Quantification represents the mean percentage of cytoplasmic area of Cellrox dye per field (n=4 wells from 3 independent experiments). **** P<0.0001, One-way ANOVA; #### P<0.001: comparison with siCont + THBP without drug treatment. **c.** Heatmaps of selected genes link to the Nrf2 pathway. qPCR screening performed on whole brain lysates isolated from P16 *Trim47+/+* and *Trim47-/-* mice exposed to normoxia condition or oxygen-induced retinopathy (OIR) model and. Data were normalized to *18S* (n=5-7 replicates per genotype and condition). # P<0.01, +/+ OIR *vs* +/+ 21%O_2_, * P<0.0001, -/- OIR *vs* +/+ OIR. One-way ANOVA. **d**. qPCR in brain ECs isolated from P16 *Trim47+/+* and *Trim47-/-* mice exposed to OIR model. Data were normalized to *Pecam1* and expressed as fold change (n=3 replicates/ genotype; 2 mice/replicate).**e.** Representative images of isolectin B4 (IB4, green or black) staining of postnatal day 16 retina from *Trim47*+/+ and *Trim47*-/- mice following OIR model characterized by oxidative stress. Right panel highlights, in blue, the avascular area of the retina after OIR. Scale bar 500 µm. **f.** Quantification represents the percentage of avascular area (µm^2^) in *Trim47*+/+ and *Trim47*-/- mice at P16 following OIR (n=9 mice per genotype). * P<0.05; Mann- Whitney test. All graphical data are mean ± s.d.

To explore the antioxidant potential of TRIM47 *in vivo*, we used mice deleted for the *Trim47* gene across all tissues and cell types (knock out (KO) mice also denoted *Trim47*-/- mice) (Supplementary Fig. 4a). *Trim47*-/- mice were viable and fertile (Supplementary Fig. 4b), developed normally, showing no significant difference in weight compared to their littermate controls (*Trim47*+/+ mice) (Supplementary Fig. 4c-e) and displaying no obvious macroscopic phenotype. The efficacy of *Trim47* deletion was confirmed at the mRNA level by qPCR on brain lysates and isolated brain ECs from *Trim47*-/- and *Trim47*+/+ mice (Fig. 3c-d and Supplementary Fig. 4f).

To challenge these KO mice at a postnatal age, we employed a pathological model of oxidative stress induced by hypoxia, namely the oxygen-induced retinopathy (OIR) model, which is characterized by increased reactive oxygen species production^40,41^. qPCR screening of *Nfe2l2* and its target genes on brain tissues from *Trim47*+/+ control mice exposed to either room air or the OIR model, confirmed that this pathological model activates the NRF2 protective antioxidant system in the central nervous system (Fig. 3c). Notably, the deletion of *Trim47* resulted in a decreased in the *Nrf2* pathway activation under both baseline and pathological (OIR) conditions (Fig. 3c). A similar reduction in Nrf2 pathway was also observed in brain ECs isolated from *Trim47*+/+ *and Trim47*-/- mice following exposure to the OIR model (Fig. 3d). Additionally, histological analysis of the retinas at p16 after exposure to oxidative stress indicated that the loss of *Trim47* was associated with an increase in avascular areas (Fig. 3e-f), suggesting that *Trim47* plays a vascular protective role in this model.

Together, these results demonstrate that TRIM47 provides antioxidant properties to brain ECs *in vitro* and protects from oxidative stress in a postnatal mouse model (OIR), at least partly by promoting the Nrf2 pathway.

### *Trim47* deletion leads to cognitive impairment

Recent large-scale genomic analyses in humans have identified *TRIM47* as a putative causal gene for common, multifactorial cSVD^21^. To investigate the impact of *Trim47* deletion on cognitive functions and brain physiology, we submitted adult *Trim47* -/- mice and their control littermates to two complementary behavioral tests assessing spatial recognition memory in the Y-maze and hippocampus-dependent spatial discrimination in the Morris water maze. Contrary to controls, male *Trim47* -/- mice (Fig. 4a-b) failed to recognize the unexplored (previously inaccessible) arm during the test phase of the Y-maze procedure, indicating an inability to form short-term spatial recognition memory. Adult *Trim47* -/- mice were also impaired in the cognitively-challenging water maze paradigm where they were unable to learn the spatial location of the hidden platform. Accordingly, latency and distance swum to the platform decreased over the 4 training days in control but not *Trim47* -/- mice (Fig. 4c-e). Potential confounders such as locomotor deficits were absent in the *Trim47* -/- mice whose swim speed was comparable to that of controls (Fig. 4f). Cognitive impairments were similar in female cohorts, suggesting that the phenotype observed in *Trim47*-deficient mice is not gender-specific (Fig. 4g-l).

**Fig. 4.**
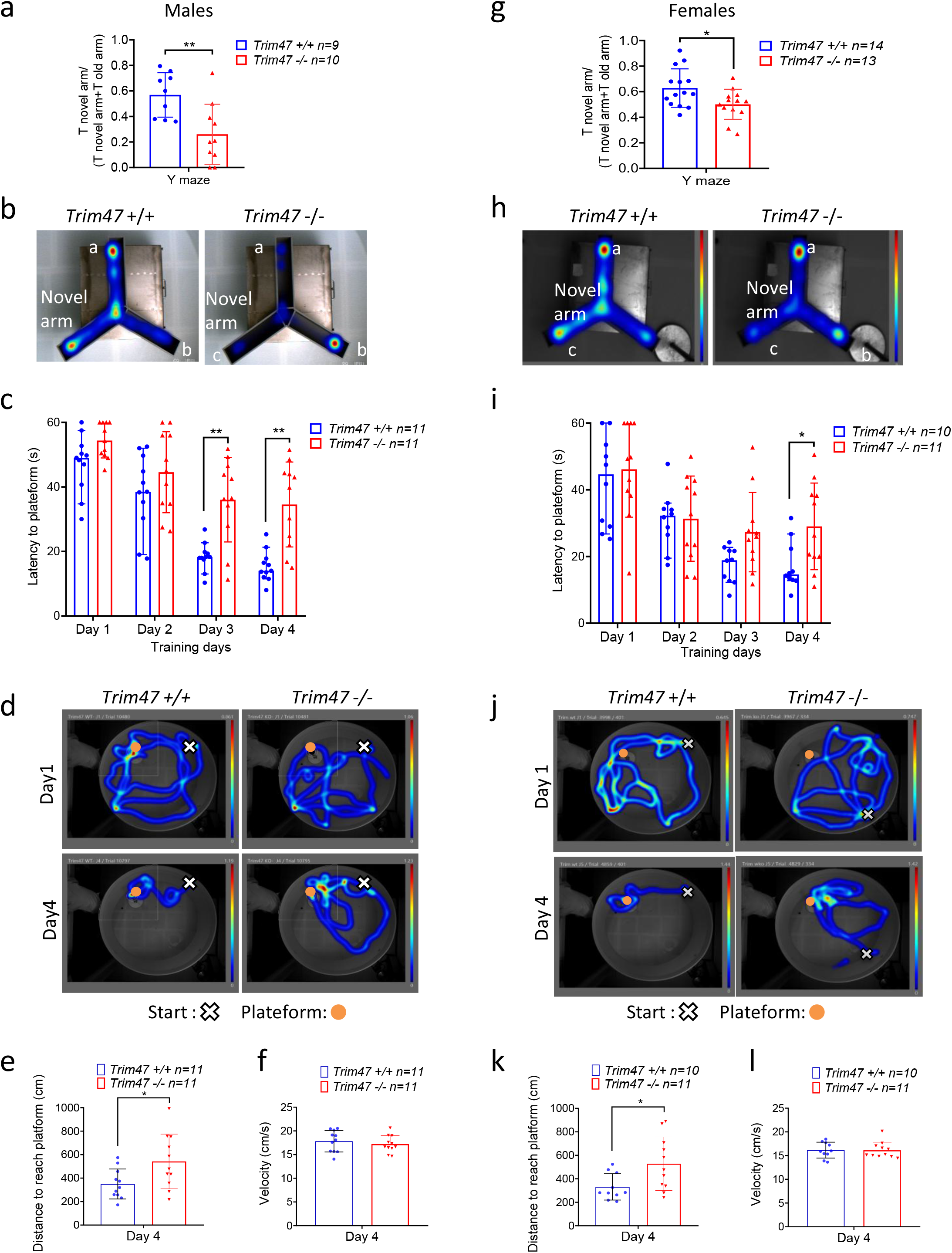
*Trim47* deletion leads to cognitive impairment **a-b.** Y-maze test was performed on males (8-9 months; *Trim47*+/+: n=9; *Trim47-/-:* n=10). Data revealed recognition memory in *Trim47*-/- mice compared to *Trim47*+/+ mice. (**a)** Y-maze data are presented as a ratio between the time spent in novel arm and the cumulative time spent in the novel and the familiar arms. **p<0.01, -/- *vs* +/+ mice, Mann-Whitney test. (**b**) Representative heatmap images of the pathes adopted in the Y-maze test. Mice started in position a; novel arm is arm c. **c-f.** Water maze test was performed in adults male (7-8 months) *Trim47*+/+ (n=11) *and Trim47*-/- (n=11) mice and revealed lower performances. *Trim47* -/- mice failed to learn the spatial position of the hidden platform (**c**) Graph shows the latency to reach the hidden platform (expressed in seconds) over 4 training days. **p<0.01 -/- *vs* +/+ mice, Two-way ANOVA with repeated measures. (**d**) Representative heatmap images of the pathes adopted by the mice during the first and last training days. The orange circle shows the position where the platform was located; the white cross indicates where the mice started the test. (**e**) Graph shows the distance to reach platform (expressed in cm) on day 4. *p<0.05, *-/- vs +/+* mice, Mann-Whitney test. (**f**) Data represent the velocity expressed as cm per second of each mouse on day 4 of water maze training. **g-h.** Y-maze was also performed in females (8-10 months; *Trim47*+/+: n=14; *Trim47-/-*: n=13). . *p<0.05, -/- *vs* +/+ mice. **i-l.** Water maze test was repeated using females (8-9 months, *Trim47*+/+: n=10; *Trim47*-/-: n=11). Female *Trim47*-/- exhibited cognitive impairments comparable to males. *p<0.05, -/- *vs* +/+ mice. All graphical data are mean ± s.d.

Interestingly, the sucrose preference test did not reveal any sign of anhedonia, a key symptom of depression in humans, suggesting that *Trim47*-/- mice develop a purely cognitive impairment without any apparent anxiety-like symptoms (Supplementary Fig. 4g).

### *Trim47* deletion triggers BBB dysfunction associated with decreased Nrf2 pathway activation *in vivo*

The enrichment of *Trim47* expression in mouse brain ECs and preliminary findings from cell-based models showing enhanced endothelial permeability following TRIM47 silencing *in vitro*^21^ pointed towards its potential role in blood brain barrier (BBB) development and/or function. Notably, Glut1 immunostaining analysis in brain sections from adult *Trim47-/-* and control mice showed no differences in vascular density in the cortical regions and hippocampus compared to control mice (Supplementary Fig. 5a-b). These data prove that *Trim47* is not essential for the development of brain blood vessels.

To determine whether the cognitive deficits observed in *Trim47* KO mice were linked to compromised BBB function, we assessed BBB integrity by injecting two different tracers. Quantification of cadaverine (Fig. 5a) and dextran (Fig. 5b) extravasation, expressed as the permeability index ratio, revealed a significantly increase in BBB leakiness in adult *Trim47*-/- mice compared to control mice. Importantly, tracer permeability in the kidney was not affected, suggesting that *Trim47* specifically acts on brain vasculature which is characterized by a specialized continuous endothelium (Fig. 5a-b). Immunostaining of brain sections with fibrinogen and podocalyxin antibodies, to visualize blood vessels, revealed the presence of fibrinogen, a blood component, outside the vessels in the hippocampus and cortical regions of *Trim47*-/- mice. In contrast, fibrinogen was confined within the vessels in control mice, confirming a localized increase in BBB permeability (in some capillaries and larger vessels) in *Trim47*-/- mice (Fig. 5c and Supplementary Fig. 5c).

**Fig. 5.**
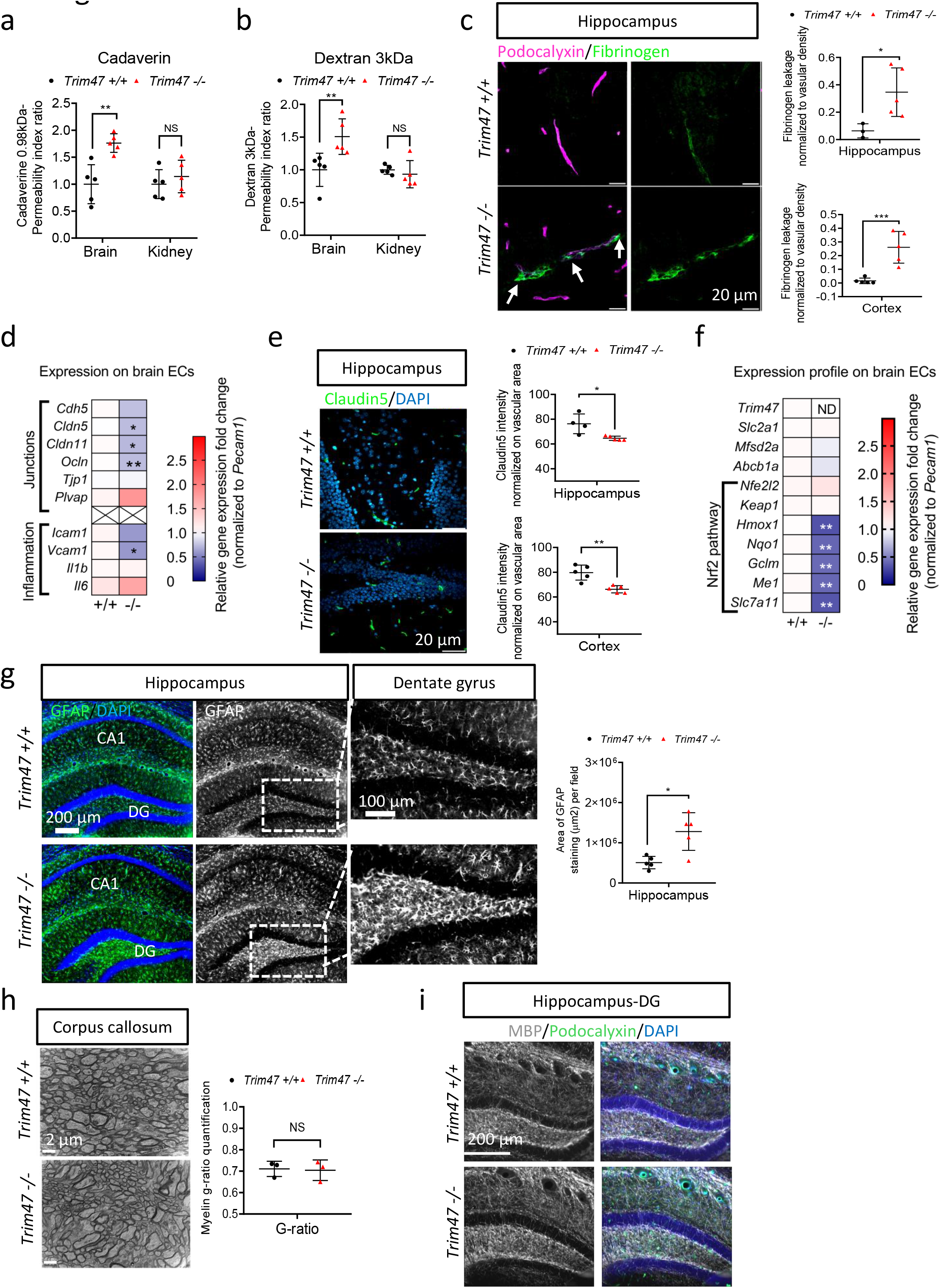
*Trim47* deletion leads to brain defects associated with Nrf2 pathway impairment **a-c**. Blood brain barrier permeability assessment was done on *Trim47+/+* and *Trim47-/-* mice (5-7 months, males). Quantification of (**a**) cadaverine and (**b**) dextran contents in brain and kidney. Data indicate the permeability index ratio (n=5 *Trim47+/+* and n=5 *Trim47-/-*). ** P<0.01, Mann-Whitney test. (**c**) Representative confocal images and quantification of fibrinogen leakage (green) in brain cryosections from *Trim47+/+* and *Trim47*-/- mice. Tissues are co-stained for podocalyxin (blood vessels, pink). Scale bars 20 µm. White arrows highlight fibrinogen extravasation in *Trim47*-/- mice. Top graph shows permeability in hippocampus (n=3 *Trim47+/+*, n=5 *Trim47*-/-) and bottom one in cortex (n=5 *Trim47+/+*, n=5 *Trim47-/-*). * P<0.05; *** P<0.001, Mann-Whitney. **d.** qPCR analysis of genes linked to junction and inflammation in brain ECs isolated from *Trim47+/+* and *Trim47-/-* mice (7-10 months). Data normalized to *Pecam1* (n=6 replicates/genotype; 2 mice/replicate, except for *Il1b*, *Il6* and *Plvap*: n=3/genotype). ND: not detectable; * P<0.05; ** P<0.01, Mann-Whitney. **e.** Representative images and quantification of claudin5 (green) in brain cryosections from *Trim47+/+* and *Trim47-/-* mice (6-7 months); nuclei identified by DAPI (blue). Scale bars 20 µm. Top graph shows claudin5 expression in hippocampus (n=4 *Trim47+/+*, n=5 *Trim47*-/-) and bottom one in cortex (n=5 per genotype). * P<0.05; ** P<0.01, Mann-Whitney. **f.** qPCR for Nrf2 target genes in brain EC from *Trim47+/+* and *Trim47*-/- mice (7-10 months). Data normalized to *Pecam1* (n=6 replicates/genotype; * P<0.05; ** P<0.01, Mann-Whitney. **g**. Representative images and quantification of GFAP (green) in thick brain coronal sections (hippocampus) from *Trim47+/+* and *Trim47-/-* mice (10 months) (n=5 *Trim47+/+* and n=5 *Trim47-/-*). Scale bars 200 or 100 µm. * P<0.05, Mann-Whitney. **h.** Electron microscopy was performed on brain thin sections to analyze white matter lesions in corpus callosum. G-ratio (ratio of the inner axonal diameter to the total outer diameter, including the myelin sheath) was measured as a readout of axonal myelination in *Trim47+/+* and *Trim47-/-* mice (8 months) (n=3 per genotype). Scale bars 2 µm. i. Representative image of MBP (white) staining in thick brain coronal sections (hippocampus) from *Trim47+/+* and *Trim47-/-* mice (8 months). Scale bars 100 µm. All graphical data are mean ± s.d.

To identify the cause of BBB defects, we analyzed the expression of key elements of tight and adherent junctions which are essential for BBB integrity, in adult *Trim4*7-/- and *Trim4*7+/+ mice. qPCR analysis of brain ECs (Fig. 5d) showed a mild but significant decrease in the expression of the tight junction components *Cldn5* (Claudin5), *Cldn11* (Claudin11), and *Ocln* (Occludin) in *Trim4*7-/- mice compared to control mice (Fig. 5d). However, the mRNA expression level of *Tjp1* (ZO1) was not affected in *Trim47*-/- mice compared to their littermate controls (Fig. 5d). Additionally, there was a trend towards a downregulation of *Cdh5* (VE-cadherin, a major element of adherens junction) in both brain ECs (Fig. 5d) and in whole brain lysates (Supplementary Fig. 5e). Importantly, immunofluorescence staining of brain sections confirmed the reduced protein levels of Claudin5 in the hippocampus and cortical regions of *Trim47*-/- mice (Fig. 5e and Supplementary Fig. 5d). These data indicated that the increased BBB permeability observed in *Trim4*7-/- mice is likely due to the decreased expression of specific tight/adherent junction components in brain ECs.

qPCR analysis confirmed the downregulation of the Nrf2 pathway activation in adult *Trim47*-deficient mice both in brain EC (Fig. 5f) and the whole brain (Supplementary Fig. 5e). This finding aligns with our *in vitro* and *in vivo* data from the hypoxia-induced oxidative stress model (OIR) at postnatal age. Blood vessels are particularly vulnerable to localized oxidative stress, and reduced antioxidant protection in ECs may further contribute to BBB impairment. Our results demonstrate that the loss of BBB integrity in *Trim47*-/- mice is associated with decreased expression of tight junction proteins. Nrf2 has been reported as a positive transcriptional regulator of Claudin5 expression in ECs^42^, suggesting that the decreased Nrf2 pathway in *Trim47*-deficient brain ECs could aggravate damage through the downregulation of Claudin5, thereby directly contributing to BBB dysfunction.

### BBB impairment-induced by *Trim47* deletion leads to brain lesions *in vivo*

Since BBB disruption can lead to the leakage and accumulation of neurotoxic material or blood components (such as fibrinogen) in the brain parenchyma, we next wanted to determine whether BBB impairment in *Trim47*-/- mice resulted in additional brain defects. Astrocytes, which are in close contact with ECs, are highly plastic and can be directly affected by BBB dysfunction and changes in their local environment^43,44^. Supporting this idea, GFAP staining indicated a significant increase in astrocyte activation in the hippocampus, particularly in the dentate gyrus region of *Trim47*-/- mice where blood vessel leakiness was observed (Fig. 5g).

As astrocytes reactivity can be modulated by neuroinflammation^43^, we next assessed the expression of adhesion molecules and cytokines in brain ECs and in whole brain lysates. qPCR analysis revealed no evidence of a pro-inflammatory endothelial state or a global pro-inflammatory profile in *Trim47*-/- mice, as shown by a reduced expression of the adhesion molecules *Icam1* and *Vcam1* (Fig. 5d and Supplementary Fig. 5e). Additionally, we analyzed immune cells and microglial markers in *Trim47*+/+ and *Trim47*-/- mouse brain samples by measuring *Cd11b* mRNA expression levels (Supplementary Fig. 6a) and performing immunofluorescence staining with Iba1 (Supplementary Fig. 6b) and Cd68 (Supplementary Fig. 6c) antibodies. Collectively, these findings indicated that *Trim47* deletion did not lead to the infiltration of immune cells or the marked activation of microglia and perivascular macrophages *in vivo*.

Since *Trim47* mutant mice show cognitive impairment, we hypothesized that the loss of *Trim47* may affect the number of neurons, axonal density and/or myelination. However, histological analyses of coronal brain sections revealed no evidence of detectable neuronal loss in *Trim47*-/- mice compared to controls, as shown by NeuN immunostaining in the hippocampus and cortical regions (Supplementary Fig. 6d-e). GWAS have identified the *TRIM47* locus as a lead genetic risk locus for high burden of WMH in humans^45^. To evaluate the impact of *Trim47* deletion on brain WM, we performed electron microscopy on brain tissues. The measurement of g-ratio^46^ in the corpus callosum region did not reveal significant axonal demyelination in *Trim47*-/- mice compared with *Trim47*+/+ mice (Fig. 5h). In line with this, myelin basic protein (MBP) staining did not show any significant reduction in myelination in the hippocampus of 10-months-old *Trim47-/-* mice (Fig. 5i), suggesting that *Trim47*-deficient mice do not display major detectable WM lesions at this stage.

### Endothelial-specific deletion of *Trim47* is sufficient to recapitulate a vascular-dementia phenotype

To demonstrate that the phenotype observed in *Trim47-/-* mice was primarily due to the loss of endothelial *Trim47*, we generated mice deleted for *Trim47* in ECs (*Trim47*^iEC-KO^) and compared them with littermate controls (*Trim47*^iEC-WT^) using an inducible CDH5-Cre line (Supplementary Fig. 7a-d). The efficiency of *Trim47* deletion was confirmed on lung ECs isolated from *Trim47*^iEC-KO^ *versus Trim47*^iEC-WT^ adult mice (Fig. 6a). Behavioral tests conducted on adult mice revealed that male *Trim47*^iEC-KO^ mice displayed impaired performance in the Y-maze (Fig. 6b-c) and water maze procedures (Fig. 6d-g) similar to the deficits observed in *Trim47-/-* mice. These finding confirm that the loss of endothelial *Trim47* is sufficient to induce cognitive impairment.

**Figure 6.**
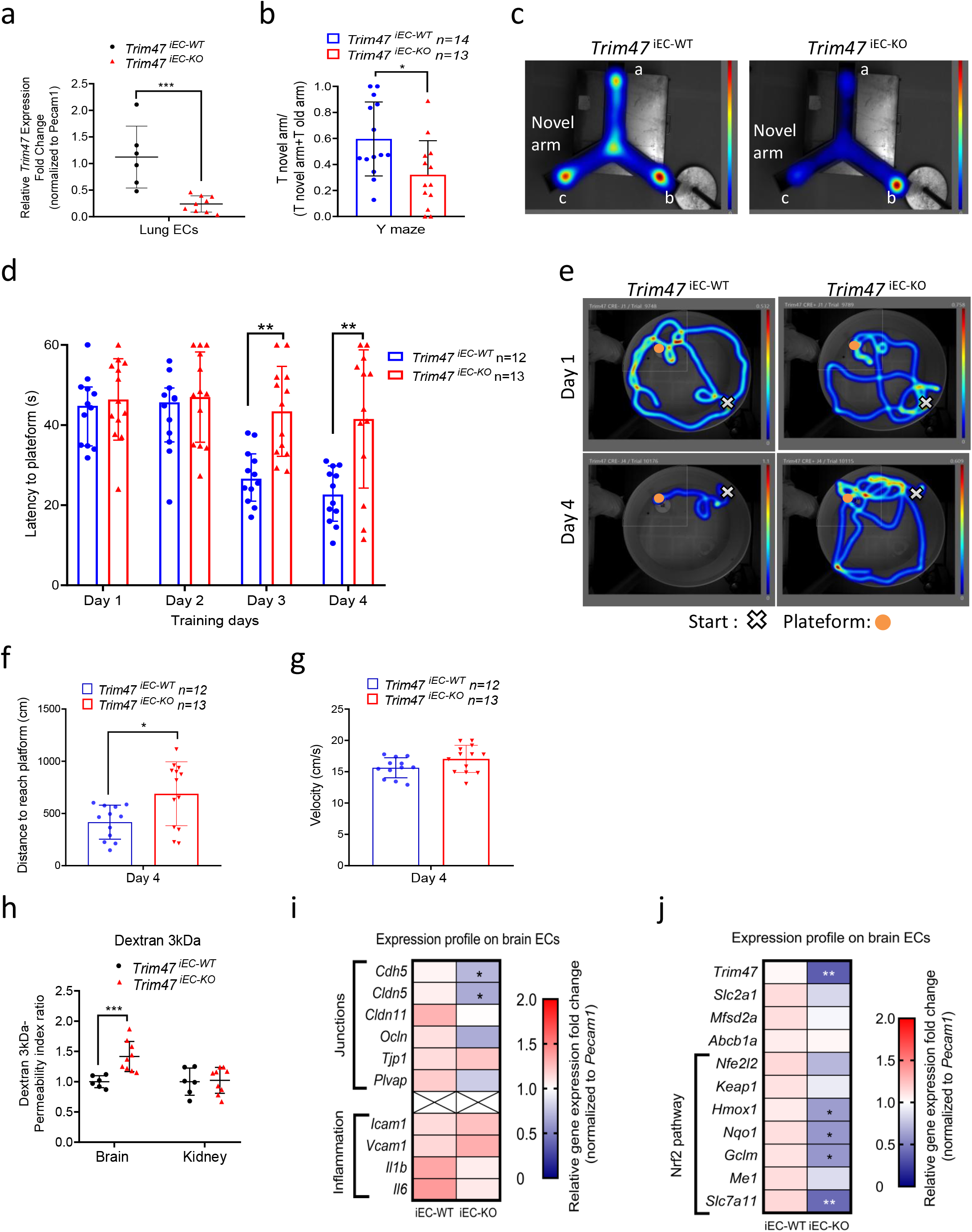
*Trim47* deletion in endothelial cells is sufficient to mimic the phenotype observed in *Trim47* full KO mice **a.** qPCR analysis for *Trim47* expression was done on lung ECs isolated from *Trim47^iEC-WT^*and *Trim47^iEC-KO^* males (5 months). Data normalized to *Pecam1* (n=6 *Trim47 ^iEC-WT^* and n=9 *Trim47 ^iEC-KO^*); *** P<0.001, Mann-Whitney. **b-c.** Y-maze test was performed on males (3-5 months; *Trim47^iEC-WT^:* n=14; *Trim47 ^iEC-KO^:* n=13). (**b**) Y-maze data are presented as a ratio between the time spent in novel arm and the cumulative time spent in the novel and the familiar arms. *p<0.05, Mann-Whitney test. (**c**) Representative heatmap images of the path for the Y-maze test. Mice started in position a; novel arm is arm c. **d-g.** Water maze test was performed on males (3-5 months; *Trim47 ^iEC-WT^:* n=12; *Trim47^iEC-KO^:* n=13). (**d**) Graph shows the latency to reach the hidden platform (expressed in seconds) for the 4 training days. **p<0.01, *iEC-WT vs iEC-KO* mice, Two-way ANOVA with repeated measures. (**e**) Representative heatmap images of the pathes for the first and last training days. (**f**) Graph shows the distance to reach platform (expressed in cm) on day 4. *p<0.05, *iEC-WT vs iEC-KO* mice, Mann- Whitney test. (**g**) Data represent the velocity expressed as cm per second (swimming speed) of each mouse on day 4 during water maze training. **h.** Assessment of BBB permeability was done on *Trim47^iEC-WT^* and *Trim47^iEC-KO^* mice (4-6 months, males). Quantification of dextran contents in brain cortex and kidney, 20 minutes after I.V injection of fluorescent tracer. Data indicate the permeability index ratio (normalization to fluorescence detected in serum) (n=5 *Trim47^iEC-WT^* and n=5 *Trim47^iEC-KO^*). * P<0.01, Mann-Whitney test. **i-j.** qPCR for (**i**) genes linked to junctions and inflammation and (**j**) Nrf2 targets in brain EC isolated from *Trim47^iEC-WT^*and *Trim47^iEC-KO^* mice (4-6 months). Data normalized to *Pecam1* (n=4-6 replicates/genotype with 2 mice/replicate; * P<0.05; ** P<0.01, ND: not detectable, Mann-Whitney. All graphical data are mean ± s.d.

Importantly, the endothelial-specific deletion of *Trim47* also led to impaired BBB function, as shown by an increased BBB permeability index following 3kDa dextran injection (Fig. 6h) and a decreased expression of key adherent and tight junction components in brain ECs isolated from these mice (Fig. 6i). Finally, qPCR analysis also highlighted the downregulation of critical Nrf2 target genes, including *Hmox1* and *Slc7a11*, in brain ECs from *Trim47*^iEC-KO^ compared to *Trim47*^iEC-WT^ controls (Fig. 6j).

These results demonstrate that the loss of *Trim47* specifically in ECs recapitulates the vascular cognitive impairment phenotype observed in the global KO mouse, underscoring the crucial role of endothelial Trim47 in maintaining brain homeostasis.

### Activation of the NRF2 pathway prevents brain defects and cognitive impairment observed in *Trim47*-deficient mice

Since we showed that TRIM47 is a key regulator of the NRF2 antioxidant pathway both *in vitro* and *in vivo*, we hypothesized that *Trim47* could promote brain homeostasis by activating the Nrf2 protective pathway in adult mice. To test this, we performed experiments on male *Trim47*+/+ and *Trim47*-/- mice, feeding them a diet containing the Nrf2 activator tBHQ (1% w/w) (Fig. 7a). As previously reported^47,48^, the tBHQ diet was well tolerated by the mice and did not induce any weight loss (Supplementary Fig. 8a). qPCR analysis confirmed that the tBHQ diet effectively prevented the impairment of the antioxidant pathway in *Trim47*-/- mice, both in brain ECs (Fig. 7b) and whole brain tissue (Supplementary Fig. 8b).

**Fig. 7.**
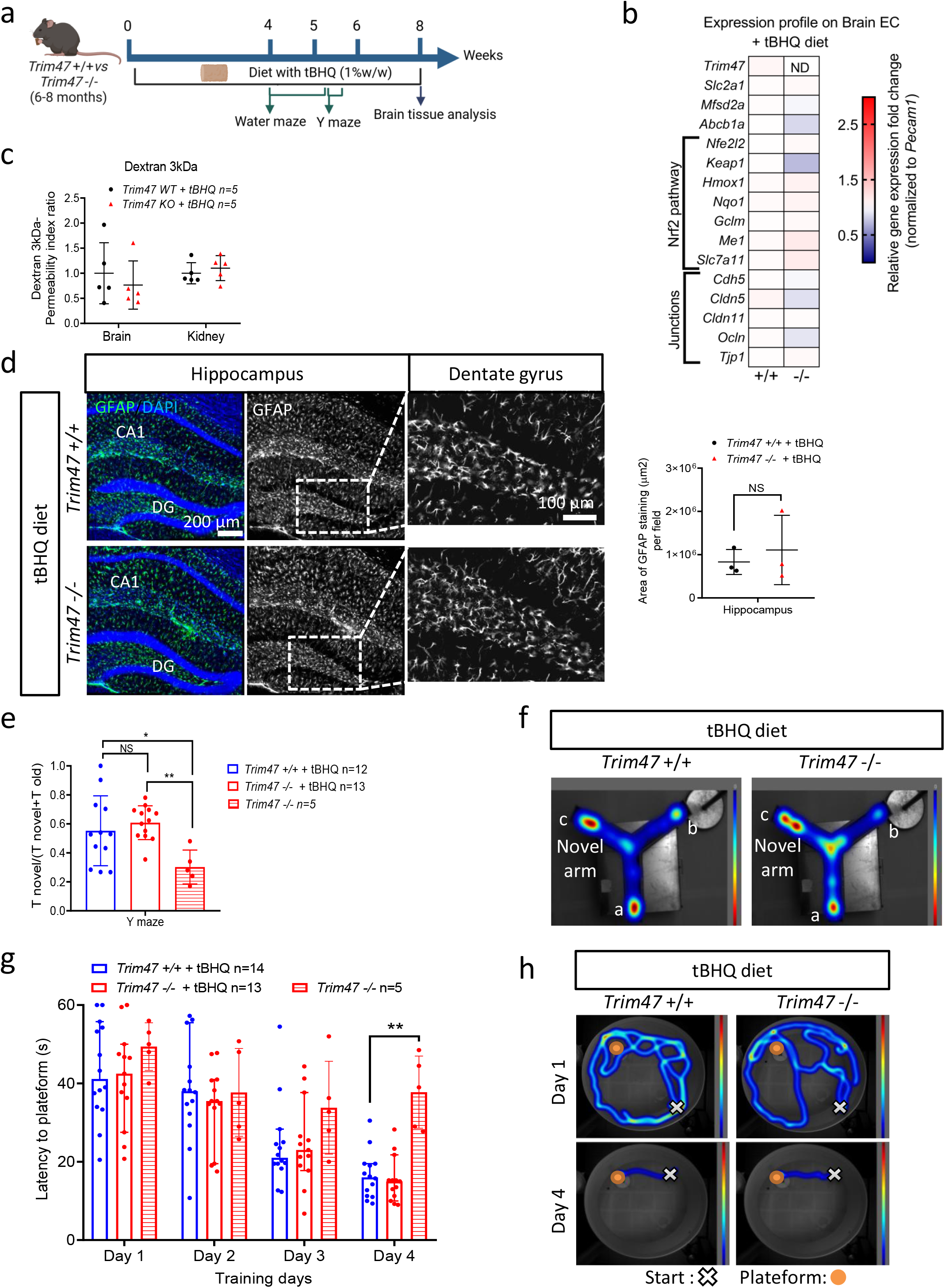
Activation of NRF2 pathway prevents cognitive impairment and brain defects observed in *Trim47* -/- mice **a.** Cartoon depicting the experimental therapeutical approach chosen in mouse. 6-8 months *Trim47+/+ and Trim47-/-* males were placed on a pellet diet containing the NRF2 pathway activator tert-butylhydroquinone (tBHQ, 1%W/W) for 1 month. Behavioral tests were then performed on these mice and on untreated *Trim47* -/- males before brain harvesting for qPCR analysis, BBB function and histology. **b**. qPCR for Nrf2 pathway performed on brain EC from *Trim47+/+ and Trim47-/-* males (8-10 months) with tBHQ diet (n=5 replicates/genotype; 2 mice/replicate). Mann-Whitney. ND: not detectable. **c**. BBB permeability assessment done on *Trim47+/+* and *Trim47*-/- males with tBHQ (8-9 months). Quantification of dextran in brain cortex and kidney (n=5 *Trim47+/+* + tBHQ and n=5 *Trim47*-/- +tBHQ). Mann-Whitney. **d.** Representative image and quantification of GFAP expression (green) in brain sections (coronal) from *Trim47+/+* and *Trim47-/-* adult mice (8-10 months) after tBHQ diet (n=3 per genotype). Scale bars 100 µm. Mann-Whitney. **e-f.** Y-maze test was performed on adult mice after a 1 month diet with tBHQ. (**e**) Data are presented as a ratio between the time spent in novel arm and the cumulative time spent in the novel and familiar arm (n=12 *Trim47+/+* + tBHQ, n=13 *Trim47-/-* + tBHQ and n=5 *Trim47-/-* without diet). *p<0.05; **p<0.01, One- way ANOVA. (**f**) Representative heatmap images of the pathes for the Y-maze test. **g-h.** Water maze test performed on adult mice after tBHQ diet. (**g**) Latency to reach platform (expressed in seconds) is shown for the 4 training days (n=14 *Trim47 +/+* + tBHQ, n=13 *Trim47 -/-* + tBHQ, n= *Trim47* -/-). *p<0.05, *Trim47 -/- vs Trim47 +/+* + tBHQ. Two-way ANOVA with repeated measures. (**h**) Representative heatmap of the pathes of *Trim47* +/+ *and Trim47* +/+ treated with tBHQ. All graphical data are mean ± s.d.

Notably, the expression of tight junction components in brain ECs was restored to normal levels in *Trim47-/-* mice treated with the Nrf2 activator compared to control mice (Fig. 7b). The diet with the antioxidant molecule prevented the increased BBB permeability (Fig. 7c), proving that antioxidant therapy successfully restored BBB integrity and function in *Trim47*-deficient mice.

In line with its known anti-inflammatory properties^49,50^, tBHQ did not normalize the mRNA levels of the adhesion molecules *Icam1* and *Vcam1* in brain ECs isolated from *Trim47* -/- mice (Supplementary Fig. 8c). Interestingly, the Nrf2 activator diet prevented astrocyte activation in *Trim47*-deficient mice, as shown by GFAP staining in the hippocampus (particularly in the dentate gyrus) (Fig. 7d).

Finally, we assessed whether increasing the Nrf2 pathway could also restore the cognitive function of *Trim47*-/- adult mice. Behavioral tests revealed that reactivation of the Nrf2 pathway *via* a tBHQ diet was sufficient to normalize the performances of the *Trim47*-/- males in both the Y-maze (Fig. 7e-f) and the water maze tests (Fig. 7g-h and Supplementary Fig. 8d-e), indicating a reversal of cognitive deficits.

Together, these data demonstrate that enhancing the Nrf2 pathway was sufficient to prevent BBB leakage, brain defects and cognitive impairment in *Trim47-/-* mice. This highlights the important role of *Trim47* in maintaining brain physiology and cognitive function, primarily through the activation of the Nrf2 antioxidant pathway.

## Discussion

In this study, we report that the cSVD risk gene *TRIM47* activates the NRF2 pathway *in vitro* and *in vivo*. TRIM47 controls brain EC resilience to oxidative stress by promoting NRF2 protein stability. *In vivo*, both global deletion and loss of endothelial *Trim47* result in a spontaneous vascular cognitive impairment phenotype characterized by increased BBB permeability, increased astrocyte activation in the hippocampus, especially the dentate gyrus and cognitive deficits in spatial recognition memory and spatial discrimination. Strikingly, activation of the NRF2 pathway using tBHQ treatment prevents these defects, showing that the protective TRIM47/NRF2 axis is required to ensure brain homeostasis.

Despite progress in understanding genetic determinants of cSVD, specific signaling pathways leading to its development are still poorly understood. Current primary prevention strategies focusing on managing blood pressure and reducing risk factors are insufficient for slowing down the progression of cSVD and preventing related stroke and VCID. There is an urgent need to develop mechanism-based therapies for cSVD, which could have a major impact at the population level for the prevention of stroke and dementia and for promoting healthier brain aging^1,9,10^. Deciphering the molecular mechanisms underlying cSVD etiology is essential for identifying novel targetable pathways. We leveraged robust genetic evidence for a causal involvement of *TRIM47* loss-of-function variants in cSVD pathogenesis, previously obtained through a combination of genomic studies in large population-based cohorts, integrated with genetically determined gene expression in vascular and brain tissues, and screening of human knock-outs in a large biobank^21^. Proof of concept *in vitro* experiments had further supported TRIM47 as a strong causal gene for cSVD^21^.

In the present study, we aim to investigate how this novel candidate selected from human studies could contribute to cSVD pathophysiology using preclinical models. We found that adult mice (males and females) deleted for *Trim47* display spatial memory deficits associated with increased BBB permeability (cadaverin, dextran, fibrinogen leakages) and decreased expression of tight junction proteins (Claudin5, Occludin) in cortical and hippocampal regions, suggesting that this pathology is driven by blood vessel damage developing at a young age. In line with this concept, endothelial-specific deletion of *Trim47* was able to fully mimic this phenotype, showing that the loss of *Trim47* in brain ECs is primarily responsible for the development of the disease. Interestingly, and supporting our findings, the genetic mouse model of endothelial Nitric Oxide Synthase (eNOS) deficiency which has been reported as a model of cerebral chronic hypoperfusion also causes spontaneous mild cognitive impairment associated with BBB leakage^51–53^.

Additionally, we describe increased GFAP-positive astrocytes in *Trim47* deleted mice specifically in the dentate gyrus and in cortical regions where blood vessel leakiness was detected. This astrogliosis in *Trim47*-deficient mice is presumably due to the localized extravasation of plasma proteins such as fibrinogen. Supporting this hypothesis, previous elegant work from other groups has demonstrated the deleterious role of fibrinogen deposits into the CNS and its numerous neuropathological effects including astrocyte and microglia activation, demyelination and neurodegeneration^54–56^. Notably, in the present work *Trim47-/-* mice do not show activation of microglia or clear signs of neuroinflammation at baseline conditions, concurring with previous studies reporting the role of *Trim47* in activating the NF-kB signalling pathway and thus playing a pro-inflammatory role in different inflammatory models^35,36^. Together, these data point toward a dual role of TRIM47, which can activate the protective NRF2 system or promote the pro-inflammatory NF-kB pathway depending on the context and the pathological model.

Our *in vitro* data identified TRIM47 as a novel and important partner of the Nrf2 antioxidant system. TRIM47 acts upstream of NRF2 and activated this pathway by inducing NRF2 protein stability, potentially through binding to its inhibitor KEAP1 in brain ECs. *In vivo*, our results confirmed TRIM47 regulation of the NRF2 system at postnatal age, in baseline and oxidative stress conditions, but also in adult mice. Crucially, we demonstrated that activation of the Nrf2 pathway with tBHQ is sufficient to prevent the BBB defects and the vascular dementia phenotype observed in *Trim47* mutant mice, proving that the endothelial TRIM47/NRF2 axis is a pivotal antioxidant mechanism actively promoting brain homeostasis.

In fact, the brain vasculature is one of the main targets of localized oxidative stress processes. Reactive oxygen species and oxidative stress have been identified as factors with a potentially strong impact on the etiology of cerebrovascular diseases^57^. Although previously underestimated, age-associated vascular oxidative stress^58^ and impairment of NRF2 signaling pathway^59^ contributing to BBB disruption^60,61^ are hypothesized to play a potential role in the pathogenesis of vascular dementia^62,63^. Nrf2 is a transcription factor that controls the expression of an array of antioxidant and detoxification enzymes^64^ and has been shown recently to regulate the expression of tight junction proteins including Claudin5, even if the exact mechanism is not fully understood^42^. Importantly, Nrf2 deficiency was shown to exacerbate obesity-^65^ or hypoperfusion-induced^66^ BBB disruption and cognitive decline in mice. Conversely, Nrf2 activation with the antioxidant sulforaphane was demonstrated to improve BBB function after ischemic stroke^67^. These data support our *in vivo* findings showing that loss of *Trim47* induces Nrf2 pathway impairment, triggering primary endothelial damages, blood vessel insults, and leading ultimately to cognitive deficits.

Taken together, our data demonstrate that *Trim47*-deficient mice display some key features of human cSVD, making this tool an innovative preclinical model of VCID that could be used by the scientific community in the vascular biology and neurosciences fields to better understand the pathophysiology of cSVD. This could be a helpful model for testing the efficacy of new therapeutical strategies developed to tackle cSVD and related VCID. As a limitation of this model, while *Trim47-/-* mice show blood vessel damages (loss of tight junction integrity, BBB leakages) associated with cognitive deficits, no major WM lesion was detectable in young and middle-aged mice (5-12 months). The addition of other risk factors such as aging (examination of 24 month-old mice) or hypertension (Angiotensin II infusion model) could increase the susceptibility of the present model to develop axonal damages or demyelination.

This work reveals a major role of TRIM47 in brain physiology and an important contribution of the TRIM47/NRF2 pathway to cSVD pathophysiology. Our findings suggest that individuals with *TRIM47* loss-of-function mutations or genetically determined lower expression of *TRIM47* may be more susceptible to developing cSVD by losing their NRF2 antioxidant protection. This highlights NRF2 targeting as a viable therapeutic strategy for cSVD and related VCID. This may pave the way to explore the use of interventions with NRF2 activators or KEAP1 inhibitors for preventing or treating cSVD and related VCID.

This study demonstrates the feasibility of using a combined bioinformatics, *in vitro*, and *in vivo* approach to functionally characterize robust genetic risk variants of cSVD. The present proof-of-concept study suggests that this strategy can efficiently inform drug discovery.

## Methods

### Cell culture and treatments

Human brain microvascular endothelial cells (HBMEC) were purchased from SicenCell (#1000) and cultured in Endothelial Cell Growth Medium-2 media (EGM-2) (Lonza). HBMEC were transfected with lentivirus or siRNA in EGM-2 media (Lonza). HeLa (ATCC CCL-2) and Hek293 (ATCC CRL-1573) were cultured in DMEM (Dulbecco’s Modified Eagle Medium, Thermo Fisher Scientific) supplemented with 10% FBS (fetal bovine serum) and 1% penicillin-streptomycin (Thermo Fisher Scientific). Cells were transfected with a pool of two siRNA against TRIM47 (siTRIM47#1 and #2, denoted as siTRIM47 in the text) (15nM for each TRIM47 siRNA, thus siRNAs at a final concentration of 30 nM) or with siRNA against NRF2 (15 nM) using INTERFERin® transfection reagent (Polyplus). Negative control siRNA (Eurogentec) was used, which is denoted as siCont. Sequences of the siRNA are listed in **Supplementary Table 1**. Plasmids were transfected using jetPRIME (Polyplus) following the manufacturer’s protocol. Lentiviruses were transduced at a multiplicity of infection of 30.

For some experiments, cells were stimulated with H2O2 (Sigma), TBHP (Luperox^®^, Sigma) or Angiotensin II (ANGII, Sigma, A9525) to induce oxidative stress conditions. In some instances, HBMEC were treated with the proteasome inhibitor MG-132 for 4 h (5 µM, M7449, Sigma) to inhibit protein degradation. For activation of the NRF2 pathway, cells were pre-treated with tert-butylhydroquinone (tBHQ) (Sigma, 112941) or dimethyl fumarate (DMF) (Santa Cruz, sc-239774).

### Plasmids and lentiviruses

The plasmid pCMV6 TRIM47 (Myc-DDK-tagged) was purchased from Origene (RC218521). It contains the full sequence of human TRIM47 (NM_033452) which was then subcloned into the lentiviral vector pRRLsin-MND-MCS-WPRE. Preparations of lentivirus encoding for human TRIM47 were produced at the Bordeaux University lentivirus platform (TBMCore).

Flag-Keap1 was a gift from Qing Zhong (Addgene plasmid #28023; http://n2t.net/addgene:28023; RRID:Addgene_28023). pCDNA3-Myc3-Nrf2 was a gift from Yue Xiong (Addgene plasmid #21555; http://n2t.net/addgene:21555; RRID:Addgene_21555).

pHOGL3/4.5 and pHOGL3/4.5 Triple mutant plasmids were a gift from Prof. Anupam Agarwal (University of Alabama at Birmingham, US). The pHOGL3/4.5 double mutant was obtained from the pHOGL3/4.5 triple mutant in which mutation C>T was created into the mutated EBox by a rapid-site-directed-mutagenesis using the following non-overlapping primers (F: cacgtgacccgccgagcata; R: ggccaggcggaacagc) and Platinum™SuperFi™II DNA Polymerase (Invitrogen). pGL3 Firefly Luciferase basic empty vector (Promega) was used as a control. pGL4.74 [hRluc/TK] vector] (Promega) Renilla luciferase vector was used as an internal control.

### Immunoblotting analysis and co-immunoprecipitation

Mouse brains were collected; cortices were dissected and then frozen in liquid nitrogen. HBMEC or mouse cortices were lysed in RIPA buffer supplemented with protease and phosphatase inhibitors using a Tissue-Lyser (Qiagen). All protein lysates were then centrifuged for 15 min at 16000 × g at 4 °C and supernatants were isolated. Proteins were loaded in 7 or 12% acrylamide gels and then transferred on a polyvinylidene difluoride (PVDF Immobilon membrane, Merck) membrane and incubated in PBS 0.1% Tween supplemented with 5% milk for 1 h to block non-specific binding. Immunoblots were next labeled with the primary antibodies listed in **Supplementary Table 2**. Primary antibodies were detected using fluorescently labeled secondary antibodies: anti-rabbit IgG Alexa Fluor™ 750 and anti-mouse IgG Alexa Fluor™ 700 (Fisher Scientific). Detection of fluorescence intensity was performed using an Odyssey imaging system (Li-COR, ScienceTec) and Odyssey ver.3 software. Fluorescence intensity was quantified by densitometry using Fiji-ImageJ software to measure protein levels and normalized against loading controls. See the source data file for the uncropped immunoblots.

For immunoprecipitation experiments, HEK293 were co-transfected with KEAP1-Flag (0.125 μg.ml−1) and TRIM47-myc-Flag (0.125 μg. ml−1). Proteins lysates were incubated with primary antibody at +4 °C, followed by incubation with Pierce™ Protein A/G Magnetic Agarose Beads (Thermo Fisher Scientific, 78609). Immunoprecipitation was carried out using primaries antibodies at 2 μg for 500 μg of protein lysates. The primary antibodies used are listed in **Supplementary Table2**.

### Reporter luciferase assay

HeLa were transfected with control or TRIM47 siRNA for 24h or transfected with either 0.1 µg of TRIM47 plasmid or pcDNA3.1 empty vector and with HO1 luciferase reporter plasmids or pGl3 vector for 24 h. Luciferase activity was measured using the Dual- Luciferase^®^ Reporter Assay System (Promega) and a Spark® Multimode Microplate Reader (Tecan). Luciferase reporter activity was normalized to the internal Renilla luciferase control and is expressed relative to the pGL3 empty vector.

### Real-time polymerase chain reaction

Total RNA from mouse tissues (brain and brain endothelial cells) and human cells (HBMEC and HeLa) was isolated by using the Direct-zol™ RNA MicroPrep kit (R2062, Zymo Research) and reverse transcribed into cDNA using the M-MLV Reverse Transcriptase (Promega). Quantitative real-time PCR was performed using the GoTaq qPCR master mix (Promega) on a QuantStudio™ 3 qPCR System (Thermo Fisher Scientific). Gene expression values were normalized to cyclophilin (PPIA) expression (human cells), *Pecam1* (mouse brain endothelial cells) or *18s* (mouse brain). See **Supplementary Tables** 3 and 4 for the list of oligonucleotides.

### Bulk RNA-sequencing on HBMEC

HBMEC were transfected with control *vs* TRIM47 siRNA for 72h. Total RNAs were isolated as described above. RNA-seq analysis was conducted on three biological replicates. Library preparation and sequencing were performed at the transcriptomic platform of the Paris Brain Institute (ICM). mRNA library preparation was performed based on the manufacturer’s recommendations (KAPA mRNA HyperPrep Kit [ROCHE]). Pooled library preparations were sequenced on NextSeq® 500 whole genome sequencing (Illumina®), corresponding to 2x30 million reads per sample after demultiplexing. The quality of raw data was evaluated with FastQC. Poor-quality sequences were trimmed or removed with Trimmomatic software to retain only good quality paired reads. Star v2.5.3a 53 was used to align reads on the human GRCh37/hg19 reference genome using standard options. Quantification of gene was done using RNA-Seq by Expectation-Maximization (RSEM) v 1.2.28, prior to normalization using the edgeR Bioconductor software package. Data analysis and visualization of the results was done with the Data Analysis Core facility from ICM and their user interface (QuBy). Differential analysis was conducted with the generalized linear model framework likelihood ratio test from edgeR using a log2FC threshold set up to 1. Multiple hypothesis adjusted p-values were calculated using the Benjamini-Hochberg procedure to control for the false discovery rate (FDR). Genes with low counts (CPM<5) were excluded from the analysis. Genes with an adjusted p-value (FDR) <0.05 were considered as differentially expressed genes (DEG). Gene set enrichment analysis was performed using the Molecular Signatures Database (MSigDB) for the Wikipathways. Volcano plots and dotplots with enrichment scores (NES) were directly downloaded from QuBy.

### Oxidative stress measurement

HBMEC seeded on 24 well plates were transfected with control or TRIM47 siRNA and then replated on Nunc™ Lab-Tek™ II Chamber Slide™ (Thermo fisher Scientific) after 24h. Cells were then pre-treated with DMSO, tBHQ (10 µM) or DMF (10 µM) for 24h and then oxidative stress was induced by TBHP (Luperox, 150 µM, 1h) or with angiotensin II (750nM, 2h). CellROX® Green reagent (C10444, Invitrogen) for oxidative stress detection and Hoechst for identifying nuclei (Thermo Fisher Scientific) were added directly to the live cells in complete EGM2 medium. After 30 minutes incubation, cells were washed 3 times in PBS and imaged with a fluorescent microscope (Observer Z.1, Zeiss). Oxidative stress was measured by the quantification of green fluorescence in cytoplasm using the “Measure nuclear and cytoplasmic intensities tool” from ImageJ macro.

### Mice and breeding

Animal experiments were performed in accordance with the guidelines from Directive 2010/63/EU of the European Parliament on the protection of animals used for testing and research and approved by the local Animal Care and Use Committee of the Bordeaux University CEEA50 (IACUC protocol #36971). General surgical procedures were conducted in mice according to ARRIVE guidelines (https://www.nc3rs.org.uk/arrive-guidelines). All animals used were retained on a C57BL/6 background and both male and female mice were used for experiments.

*Trim47* full KO mice (C57BL/6N-Trim47em1(IMPC)Bay/Mmmh) were purchased from the national public repository system for mutant mice Mutant Mouse Resource and Research Center. The inducible endothelial-specific *Trim47* knockout mouse model (*Trim47*iEC-KO) was generated by breeding *Trim47 ^fl/fl^* mice (strain ID: T005408, GemPharmtech) with Cdh5(PAC)-CreERT2 mice^68^. Endothelial deletion of *Trim47* was induced in adult mice, by tamoxifen injection (5648, Sigma) (five injections of 0.5 mg daily). All experiments were conducted using littermate controls denoted in the text as *Trim47* +/+ for the full KO mice and as *Trim47*^iEC-WT^ for the endothelial-specific deleted mice line.

For the activation of the Nrf2 pathway, adult *Trim47* -/- and *Trim47* +/+ males were kept on a diet containing tBHQ (Sigma, 1% w/w). The food pellets were supplemented with tBHQ by the manufacturer Safe. Body weight and food intake were monitored regularly. Behavioral tests were performed after a 1 month-diet and tissues were collected for histology, BBB permeability assessment or transcriptomic analysis after 2 months.

### *In vivo* model of oxidative stress - oxygen-induced retinopathy model

The oxygen-induced retinopathy model (OIR) was carried out as described previously^69,70^. Mice pups and their nursing mother were exposed to 75% oxygen for 5 days from age P8 to age P13 in a hyperoxic chamber (ProOx Model 110, BioSpherix). At P13, they were returned to normal room air. Pups were euthanized at P16 to assess the degree of vascular regression (avascular area) and to evaluate retinal revascularization with pathological vessels (NV tufts). Only the pups weighing >4.5 g at P16 were included in the study since low weight impact the outcome and severity of the OIR model^71^. Mouse eyes at P16 postnatal stage were dissected and stained with isolectin-B4 in PBS supplemented with 3% bovine serum albumin, 1% donkey serum and 0.5% Triton. Retinal vasculatures were imaged with a fluorescent microscope (Observer Z.1, Zeiss). Avascular retinal areas were quantified with Zen blue software (version 3.5, Zeiss), normalized to the total retinal area and expressed as a percentage compared to littermate controls.

### Behavioral tests

#### Y-maze

The Y-maze apparatus consisted of three arms (8 × 30 × 15 cm) separated by an angle of 120°. The recognition memory procedure took advantage of the innate tendency of rodents to explore novel environments. It consisted of four trials performed over 2 consecutive days. During the exploration (encoding) trial, mice were allowed to explore only two arms (starting arm and familiar arm) of the maze for 5 minutes, with the third arm being blocked (novel arm). The mice were given two exploration trials the first day with an inter-trial interval of 3 h and one additional trial the second day to reinforce encoding. Three hours after the last exploration trial, mice underwent a recognition trial. Each mouse was put back in the starting arm of the maze, with free access to all three arms for 5 min. Time spent by mice in each arm and their trajectories were recorded and analyzed using an automated videotracking system (EthoVisionXT 16, Noldus). Discriminating the novel arm from the two familiar arms was used as an index of spatial recognition memory. Memory performance was expressed as the percentage of time spent in novel arm calculated as follows: (time spent in novel arm/time spent in all three arms) × 100.

#### Morris water maze

This test used a circular pool (180 cm in diameter) located in a dimly lit room decorated with various distal cues. The pool contained opaque water (22 ± 1 °C) and was subdivided into four virtual quadrants (A, B, C and D). A circular escape platform (10 cm in diameter) was submerged 1 cm beneath the water surface. The hidden platform remained in the same fixed position (quadrant D) during training. For each training trial, mice could swim for a maximum of 60 seconds to locate the hidden platform and were allowed to stay on it for 15 seconds upon finding it. Mice unable to locate the platform were gently guided to it. Mice were trained over 4 consecutive days and given 2 daily trials separated by an inter-trial interval of 3 h. The performance of each mouse (latency to reach the platform, distance swum, swim speed and trajectory) was acquired and analyzed using an automated videotracking system using (EthoVisionXT 16, Noldus).

#### Sucrose preference test

To assess anhedonia and anxiety, we performed a sucrose preference test on adult mice. The mice were previously accustomed to having 2 bottles in their cage, one week before the test. Mice were then offered two 50-ml bottles filled with water and 1% sucrose for 4 days. The 1% sucrose solution was then replaced by a 3% sucrose solution which was again replaced 4 days later by a 9% sucrose solution. Each day, water and sucrose amounts were monitored by weighing the water bottles, and the positions of the bottles in the cages were counterbalanced to avoid spatial preference. Sucrose preference was determined as the percentage of sucrose intake normalized to the total liquid intake.

### Endothelial cells isolation

#### Brain endothelial cells

Mouse brains collected from 2 adult mice were pooled and transferred into a petri dish containing cold HBSS without calcium/magnesium. Cerebellum and brain stem were then removed, leaving cerebral hemispheres intact. Each cerebral hemisphere was then cut in 4 sagittal slides before being enzymatically and mechanically digested using a gentleMACS^TM^ Dissociator (#130-093-235, Miltenyi Biotec, 37C_ABDK_01 program) and the Adult Brain Dissociation kit (#130-107-677, Miltenyi Biotec) following the manufacturer. Brain lysates were filtered through a 70 µm MACS Smart Strainer (#130-198-462, Miltenyi Biotec). Incubation of the filtered suspension with Myelin Removal Beads II for 15 minutes (#130- 096-733, Miltenyi Biotec) allowed depletion of myelin. ECs enrichment was performed by incubating the suspension with CD31 Microbeads for 30 minutes (#130-097-418, Miltenyi Biotec). The positive selection of ECs was done using LS column (#130-042-201, Miltenyi Biotec) placed on a magnetic separator (MACS® MultiStand and QuadroMACS^TM^ Separator, Miltenyi Biotec). Endothelial cells were centrifuged (10 minutes, 500 g) and cell pellet was resuspended in 300 µl of TRI Reagent® (MRC gene) and then processed for RNA isolation.

#### Lung endothelial cells

Mouse lungs were collected from adult *Trim47* or *Trim47* -/- mice, minced with a scalpel and then digested using a gentleMACS^TM^ (37C_m_LDK_1 program) and the Lung Dissociation kit (P) (Miltenyi Biotec, Cat#130-095-927). Lung lysates were filtered through a 70 µm MACS Smart Strainer. Endothelial cell enrichment was performed by incubating the suspension with CD31 Microbeads for 15 minutes (#130-097-418, Miltenyi Biotec). Endothelial cells were centrifuged (10 minutes, 500 g) and cell pellet was resuspended in TRI Reagent® and then processed for RNA isolation.

### Blood brain barrier permeability assessment *in vivo*

To assess BBB permeability, fluorescent tracers were injected intravenously (retro-orbital) in adult (5-6 months) mice and left to circulate for 20 min as described previously^72^. Cadaverine (0.95 kDa) conjugated to Alexa Fluor-555 (A30677, Invitrogen) was injected at a concentration of 100 µg Cadaverine/20 g of mice and lysine-fixable 3 kDa dextran conjugated to tetramethylrhodamine (D3308, Invitrogen) at a concentration of 250 µg dextran/20 g. To assess tracers leak, a 300 µL blood sample was collected by cardiac puncture and centrifuged at 10 000 g, 10 min at 4 °C for serum preparation. Animals were then perfused in the left ventricle with warmed PBS. One hemi-brain (free of olfactory lobes and cerebellum) and kidney were then collected, and their weight measured before being snap-frozen in liquid nitrogen. Tissues were homogenized in PBS using a Tissue Lyser (Qiagen) and centrifuged at 15,000 g, 20 min, and 4 °C. Fluorescence measurement (RFUs) of 50 μL of diluted serum, brain and kidney supernatants was done using a Spark® Multimode Microplate Reader (Tecan). A Permeability Index (PI) was calculated as follows: PI (mL/g) = (Tissue RFUs/g tissue weight)/(Serum RFUs/mL serum) for each animal. The PI of each animal was divided with the mean PI of the *Trim47* +/+ group and data were represented as a ratio of PI. For additional histological analysis, the contralateral hemi-brain was directly OCT-embedded and preserved for immunohistochemistry.

### Immunofluorescence analysis of brain tissues

10 µm cryosections (sagittal) were prepared using a cryostat, then fixed with 4% PFA for 10 min at room temperature, blocked with blocking solution (PBS1X, 1 % BSA, 0.5 % Triton) for 1 h and incubated for 1.5 h at room temperature with primary antibodies diluted in buffer containing PBS1X, 0.5 % BSA and 0.25 % Triton X. For other histological analysis, brains were collected and placed in 4% PFA overnight at 4 °C. Brains were then washed 2 times 1h with PBS1X and embedded in 4% agarose. 50 µm coronal sections were prepared using a Leica VT 1000S vibratome and placed in storage solution (Glycerol 30%, Ethylene glycol 30%, PBS-1X 40%) -20 °C. Brain coronal sections were blocked with PBS 1X, 10% donkey serum, 0.05% Triton)) for 4h prior to incubation with primary antibodies for 48h and then with secondary antibodies (1/400 dilution). After 3x 30 min wash with PBS1X, sections were mounted using mounting medium (Fluoromount G with DAPI, #00-4958-02, Thermo Fisher Scientific).

Sections were imaged with a fluorescent microscope (Observer Z.1, Zeiss) and with a confocal microscope (LSM 900, Zeiss). Images were analyzed with Zen blue (version 3.5, Zeiss) and ImageJ (version 2.3.0, NIH).

### Transmission electron microscopy

Animals were perfused in the left ventricle with 0,9% NaCl and 20 IU/mL heparin, then 4% PFA with 2.5% glutaraldéhyde in 0.1M Phosphate Buffer pH7.4, and 4% PFA in 0.1M Phosphate Buffer pH7.4. Brains were collected and 150 µm coronal sections were prepared using a vibratome and placed in 4% PFA overnight at 4 °C. Samples were post-fixed with 1% osmium tetroxide and dehydrated with graded ethanol series. A final dehydration step is performed with acetone and embedded in Epon. Ultrathin sections (70 nm) were cut by using an ultramicrotome (Leica EM UC6, Leica Microsystems) and observed under a transmission electron microscope (MET 120kV HT7650, HITACHI, Japan).

### Statistical analysis

All *in vitro* results presented in this study are representative of at least three independent experiments. ‘*n*’ represents the number of biological replicates unless otherwise stated. All *in vivo* experiments were done on littermates with similar body weights per condition. Males and females were analyzed separately for behavioral tests. Data are shown as the mean ± or standard deviation (s.d) as stated in the text. Statistical analysis was performed using GraphPad Prism 8 software. A two-sided unpaired *t-test* was performed for statistical analysis of two groups for *in vitro* data. A two-sided Mann–Whitney test was performed for the analysis of two groups for *in vivo* results. One-way ANOVA followed by Bonferroni’s multiple comparisons test was performed for statistical analysis between 3 or more groups.

Two-way ANOVA with multiple comparisons test was performed to evaluate mouse performances (2 or 3 groups) and progression over time during the Water maze. Differences were considered statistically significant with a P value < 0.05.

## Data availability Statement

The raw reads for bulk RNA-sequencing in FASTQ format have been deposited in Gene Expression Omnibus (GEO) database under accession number: GSE279052. All other data supporting the findings of this work are available within the paper and its Supplementary Information file. Uncropped immunoblots are presented in Supplementary data 5. Any other data are available from the corresponding author upon reasonable request.

## Acknowledgment

We thank Professor Anupam Agarwal (The University of Alabama at Birmingham, School of Medicine, United States) for providing the HO1 promoter luciferase constructs. We thank Philippe Alzieu, Sylvain Grolleau and Maxime David for their technical support and Christelle Boulle for administrative assistance. This work benefited from equipment and services from the Bordeaux Imaging Center (BIC) platform and the iGenSeq (RNA sequencing) and iCONICS (RNAseq analysis) core facili es at the ICM (Ins tut du Cerveau et de la Moelle pini re, PARIS, France).

## Funding

This project is supported by a grant overseen by the French National Research Agency (ANR) as part of the Investment for the Future Programme ANR-18-RHUS-0002 (SHIVA Project) and by the Precision and Global Vascular Brain Health Institute (VBHI) funded by the France 2030 IHU3 initiative.

## Author contribution

C.D. contributed to study design, performed experiments, analyzed and conceptualized results, V.D., R.B., S.R, M.B., C.C. and C.Proust performed *in vivo* experiments, JV and BJV contributed to experimental design and performed *in vitro* experiments and histology, J.L.M and B.B provided equipment for behavioral tests, analyzed data for behavior assessment, A.M. contributed to scientific discussion, S.D contributed to scientific discussion and manuscript writing, T.C. designed the study, conceptualized results, and wrote the manuscript, C.Peghaire designed the study, performed experiments, analyzed and conceptualized results and wrote the manuscript.

## Competing interests

The authors declare no competing interests.

## Notes

https://www.ncbi.nlm.nih.gov/geo/query/acc.cgi?acc=GSE279052

### Competing Interest Statement

The authors have declared no competing interest.

### Summary of Updates

1) Abstract, introduction and discussion section: rewritting of the manuscript and addition of relevant references to emphasize the rational, the clinical impact and relevance of the paper 2) Data availability Statement: a GSE number was added for the RNAseq data released and deposited on GEO 3) Supplementary data: uncropped immunoblots are presented in Supplementary data 5.

