## Supplementary information for "Cerebral Small Vessel Disease genetic determinant TRIM47 controls brain homeostasis *via* the NRF2 antioxidant system"

TRIM47 - NRF2 pathway and SVD

Title:

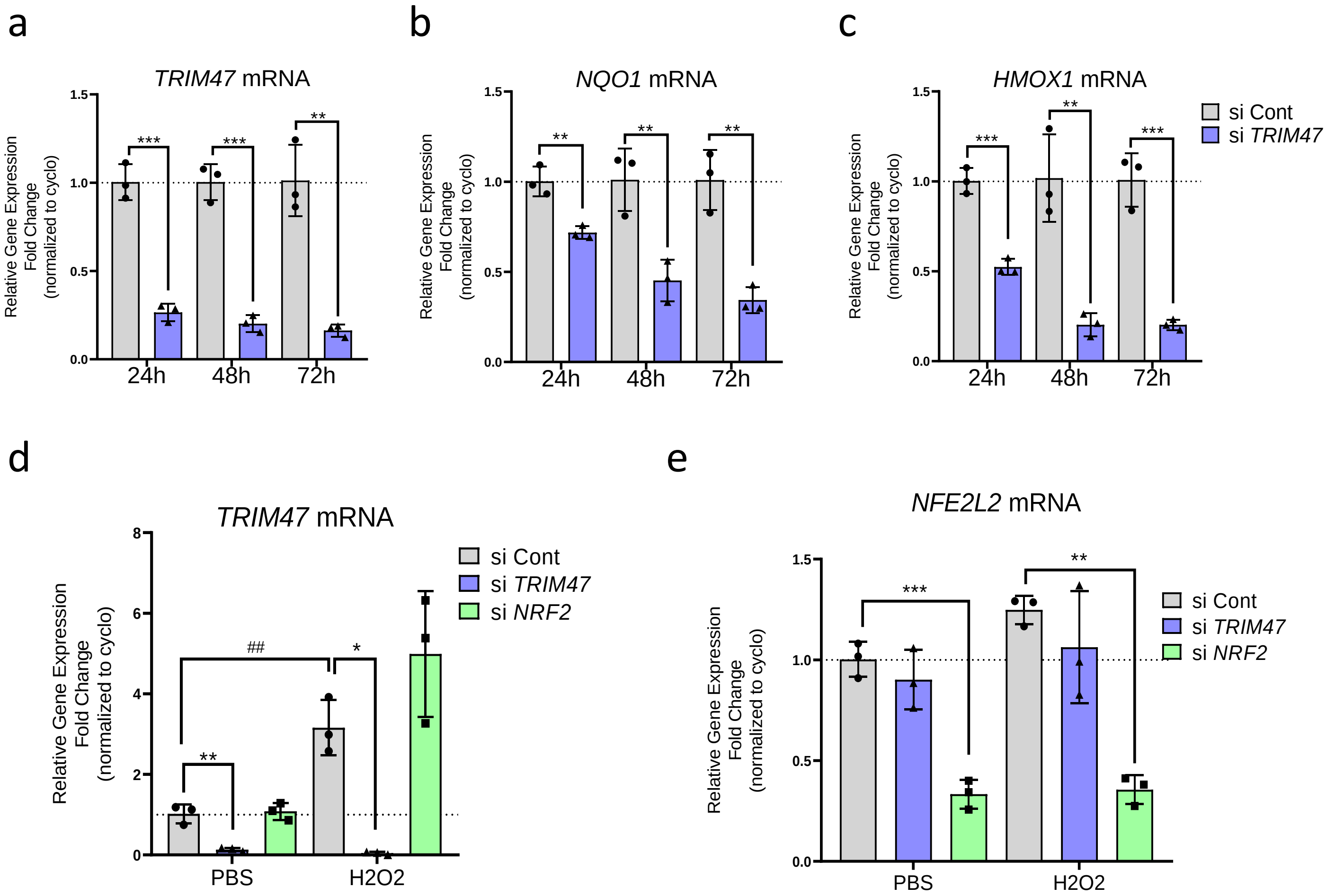


**Supplementary Figure 1**: Regulation of NRF2 target genes by TRIM47 in human brain microvascular endothelial cells (HBMEC). **a-c**. qPCR analysis of (**a**) *TRIM47*, (**b**) *NQO1* and (**c**) *HMOX1* expression in control (siCont) and *TRIM47*-deficient (si*TRIM47*) HBMEC after 24, 48 and 72 hours siRNA treatment. Data were normalized to cyclophilin (n=3 experiments). **: $p<0.01$; ***: $p<0.001$, Student's *t*-test. **d-e**. qPCR analysis of (**d**) *TRIM47* and (**f**) *NFE2L2* (NRF2) gene expression in control, *TRIM47* or *NRF2* siRNA-treated HBMEC for 72h and treated with either PBS or H2O2 (200 uM for 3h). Data were normalized to *cyclophilin* (n=3 experiments). * $P<0.05$; ** $P<0.01$; *** $P<0.001$, One-way ANOVA, ## $P<0.01$, Student's *t*-test. All Graphical data are mean ± s.d.

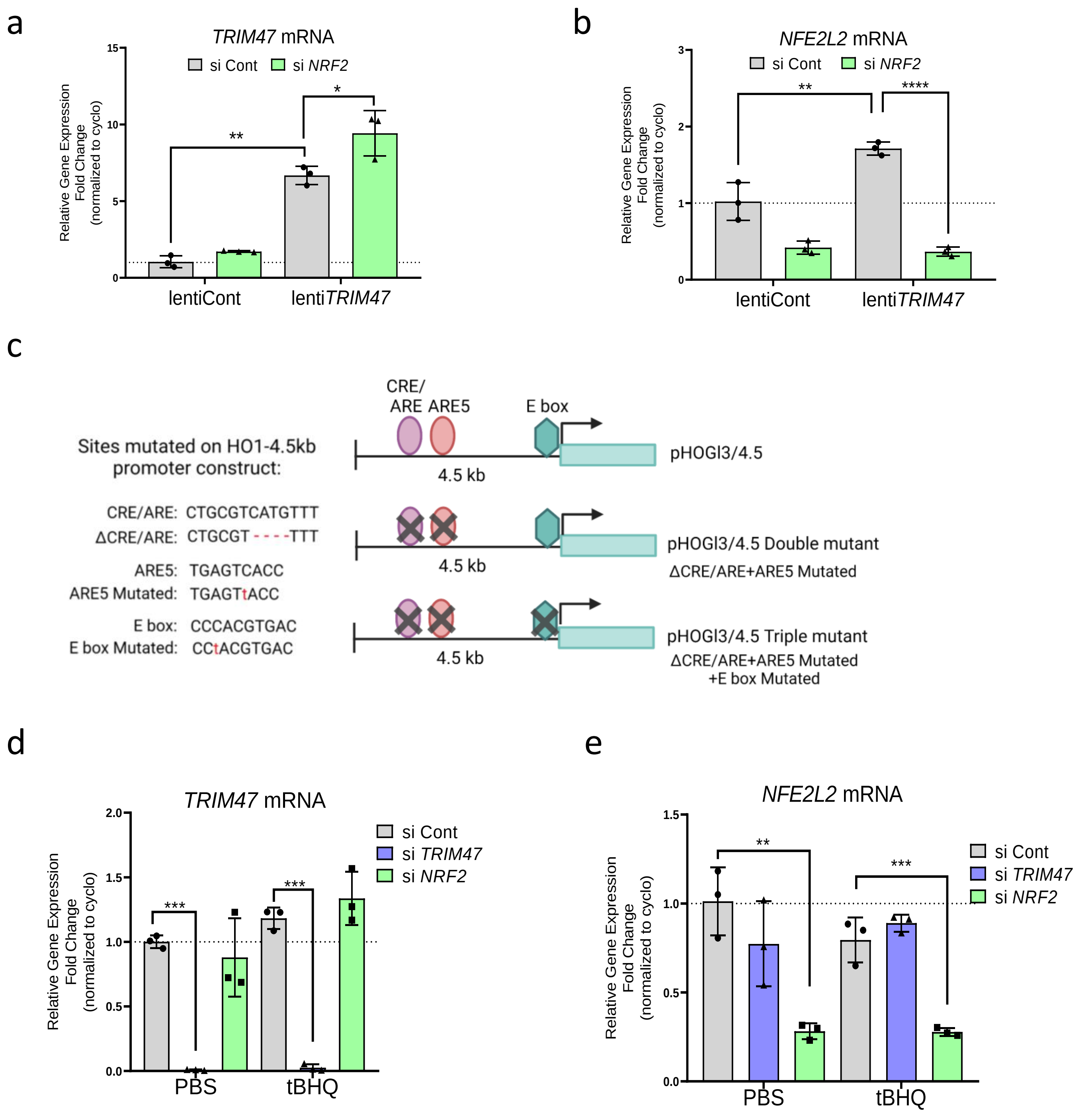


**Supplementary Figure 2**: Controls and tools for the experiments showing TRIM47 cooperation with the transcription factor NRF2 in HBMEC. **a-b**. qPCR analysis of (**a**) *TRIM47* and (**b**) *NFE2L2* expression in siCont or si*NRF2* treated HBMEC and co-transduced with a control or TRIM47 lentivirus for 24h. Data were normalized to *cyclophilin* (n=3 experiments). ** P<0.01; *** P<0.001; One-way ANOVA. **c.** Cartoon depicted the HO1 tools used for promoter luciferase reporter assay in HeLa. Wild type HO1 promoter-luciferase construct (HO1-4.5bp), construct mutated for CRE/ARE and ARE5 sites (HO1-4.5bp double mutant) and construct mutated for CRE/ARE, ARE5 sites and E box (HO1-4.5bp triple mutant, used as a control for the experiment). **d-e**. qPCR analysis of (**d**) *TRIM47* and (**e**) *NFE2L2* expression in HBMEC treated with control, *TRIM47* or *NRF2* siRNA for 72h and with tBHQ for 24h (n=3 experiments). ** P<0.01; *** P<0.001; One-way ANOVA. All Graphical data are mean ± s.d.

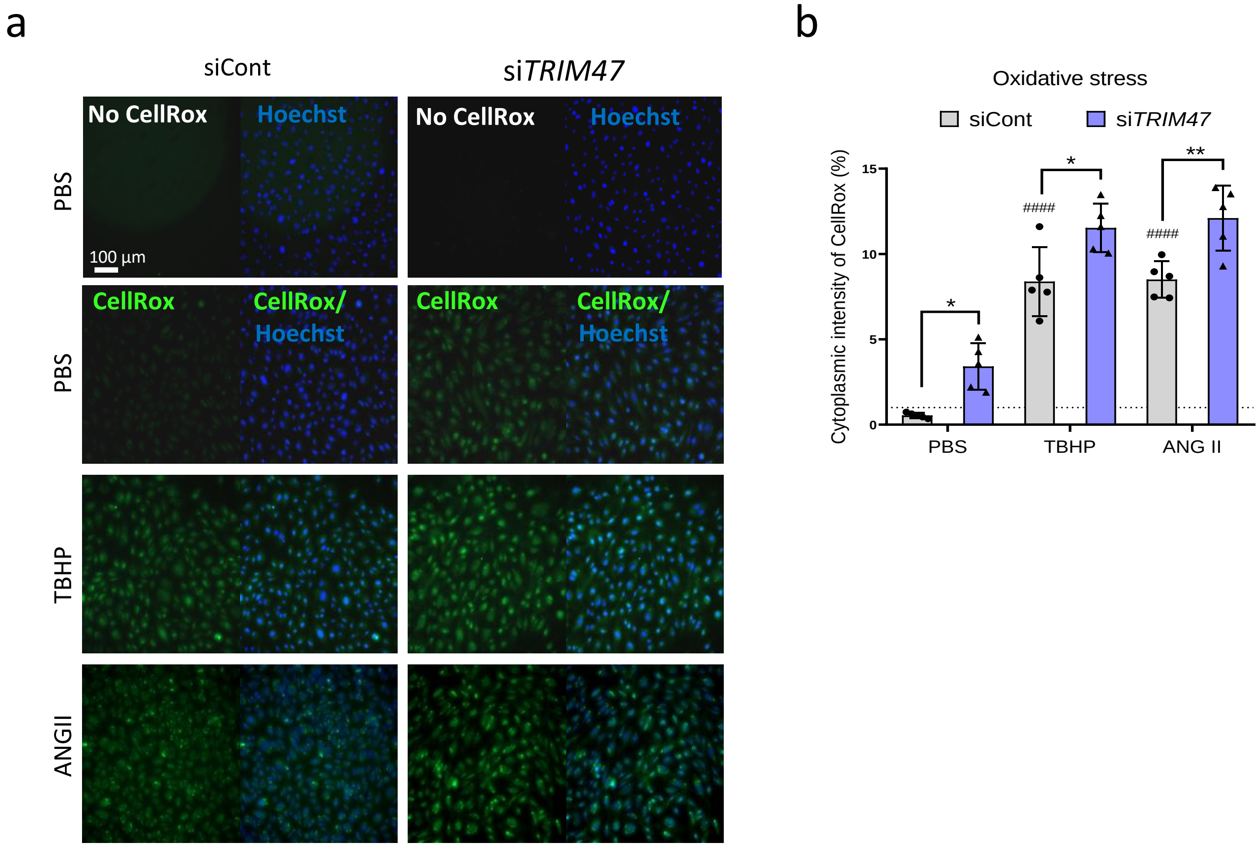


**Supplementary Figure 3**: TRIM47 displays antioxidant properties in HBMEC.

**a-b.** Representative immunofluorescence image (**a**) and quantification (**b**) of CellRox dye (green) in HBMEC transfected with siCont or si*TRIM47* for 24 h and treated with PBS or tert-butyl hydroperoxide (TBHP, 200 µM for 1 h) or Angiotensin II (ANGII, 500 nM for 2h); nuclei are identified by Hoechst (blue). Scale bar 100 µm. Quantification represents the mean percentage of cytoplasmic area of Cellrox dye per field (n=5 wells from 3 independent experiments). * P<0.05, ** P<0.01, One-way ANOVA; #### P<0.001: compared to siCont + THBP without drug treatment. Graphical data are mean ± s.d.

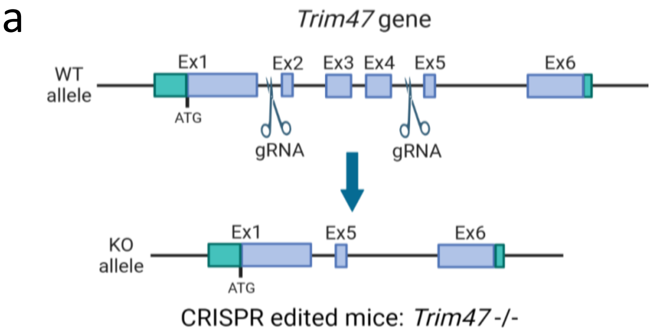


b

| genotype | +/+ | -/- | Total |
|---|---|---|---|
| % observed | 23 | 22 | 100 |
| % -mendelian ratio | 25 | 25 | 100 |

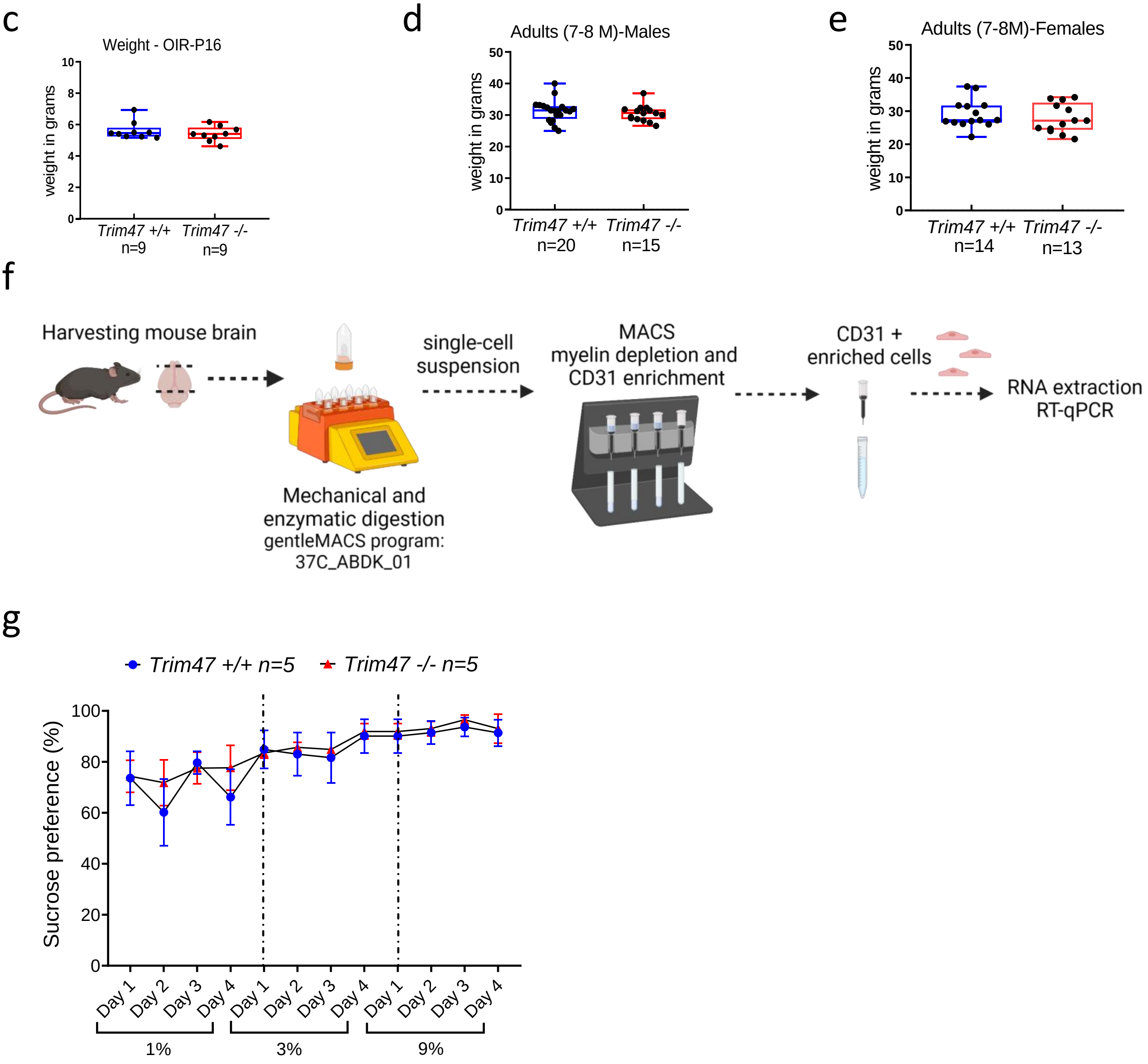


**Supplementary Figure 4**: Generation and characterization of mice deleted for *Trim47* in all tissues. **a.** Cartoon depicting the strategy for generating mice globally deleted for *Trim47* using CRISPR deletion of exons 2 to 4. **b.** Expected Mendelian ratio and percentage of mice observed with +/+ or -/- genotype is reported in the table. **c-e.** Body weight (in grams) of (**c**) postnatal day 16 mice after OIR model, adult (**d**) males and (**e**) females +/+ and -/- mice. **f.** Workflow of mouse brain endothelial cells isolation protocol using gentle MACS for mechanical and enzymatic digestion, then myelin removal step and enrichment with CD31 magnetic beads. **g.** Sucrose preference test was performed to asses anhedonia and anxiety in adults +/+ and -/- mice (males, 8 months, n=5 mice per genotype) and is expressed as the percentage of sucrose intake normalized to the total amount of liquid (water and sucrose) intake. Graphical data are mean ± s.d.

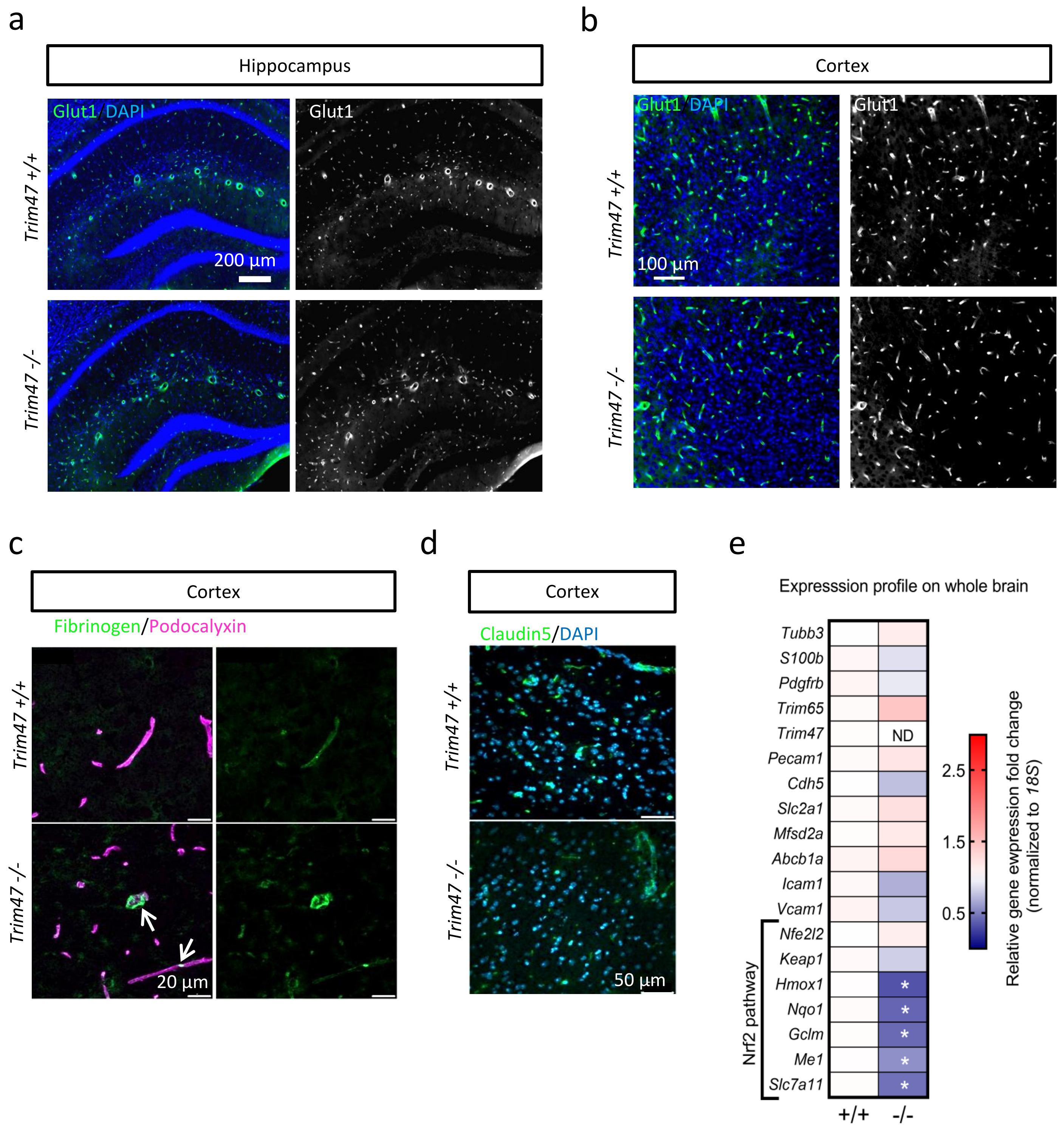


**Supplementary Figure 5**: Histological and transcriptomic analysis of *Trim47+/+* and *Trim47-/-* adults mice. **a-b.** Representative images of Glut1 (green or grayscale) in brain sections (coronal, 50 um) from *Trim47*+/+ and *Trim47*-/- mice showing blood vessels in (**a**) hippocampus and (**b**) cortical regions. Nuclei are stained with DAPI (blue). Scale bars 200 or 100 µm, as noted. **c**. Representative confocal images of fibrinogen leakage (green) in brain cryosections from *Trim47*+/+ and *Trim47*-/- mice. Tissues are co-stained for podocalyxin (blood vessels, pink). Scale bars 20 µm. White arrows highlight fibrinogen extravasation in *Trim47-/-* mice. **d.** Representative images (cryosections) of claudin5 (green) in cortical regions of *Trim47+/+* and *Trim47-/-* mice (6-7 months); nuclei identified by DAPI (blue). Scale bars 50 µm. **e.** qPCR screening of whole brain from *Trim47*+/+ and *Trim47*-/- mice (7-10 months). Data normalized to *18S* (n=4 *Trim47*+/+, n=4 *Trim47* /-). ND: not detectable; * P<0.05, Mann-Whitney.

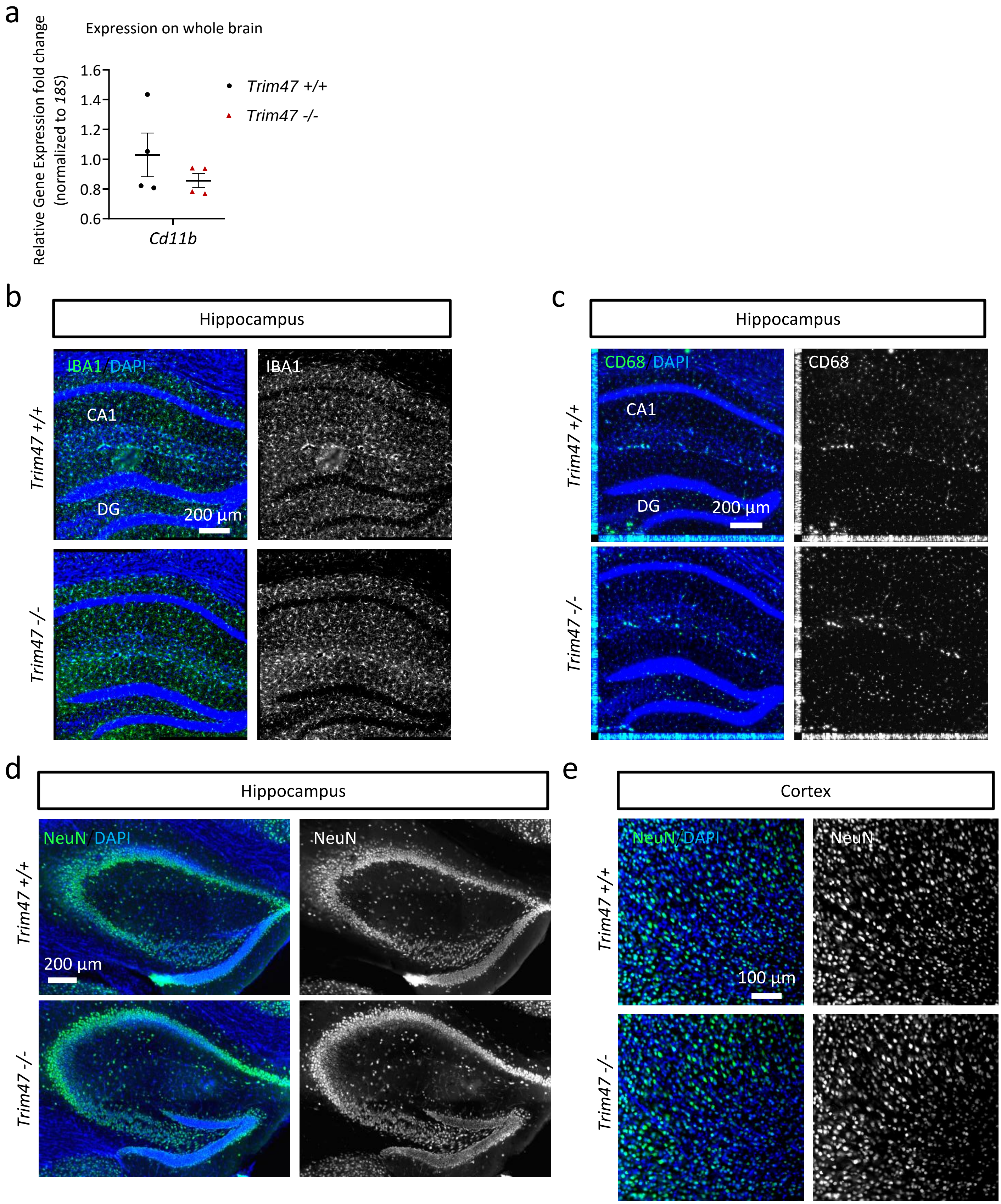


**Supplementary Figure 6**: Effects of *Trim47* deletion on non-endothelial cells in adult mouse brain.
**a.** mRNA expression profile of the microglia marker *Cd11b* in whole brain lysates from *Trim47+/+* and *Trim47-/-* mice (7-10 months). Data normalized to *18S* (n=4 *Trim47+/+,* n=4 *Trim47-/-*). **b-c**. Representative images of (**b**) Iba1 (microglia) (green/grayscale) and (**c**) Cd68 (macrophages) (green/grayscale) immunostaining in brain sections (coronal, 50 um, hippocampus) from *Trim47*+/+ and *Trim47-/-* mice. Scale bars 200 µm. CA1: cornu ammonis region 1, DG: dentate gyrus **d-e**. Representative images of Neun staining (neurons) (green/grayscale) in brain cryosections from *Trim47*+/+ and *Trim47-/-* mice in (**d**) hippocampus and in (**e**) cortex region; nuclei identified by DAPI (blue). Scale bars 200 or 100 µm. All Graphical data are mean ± s.d.

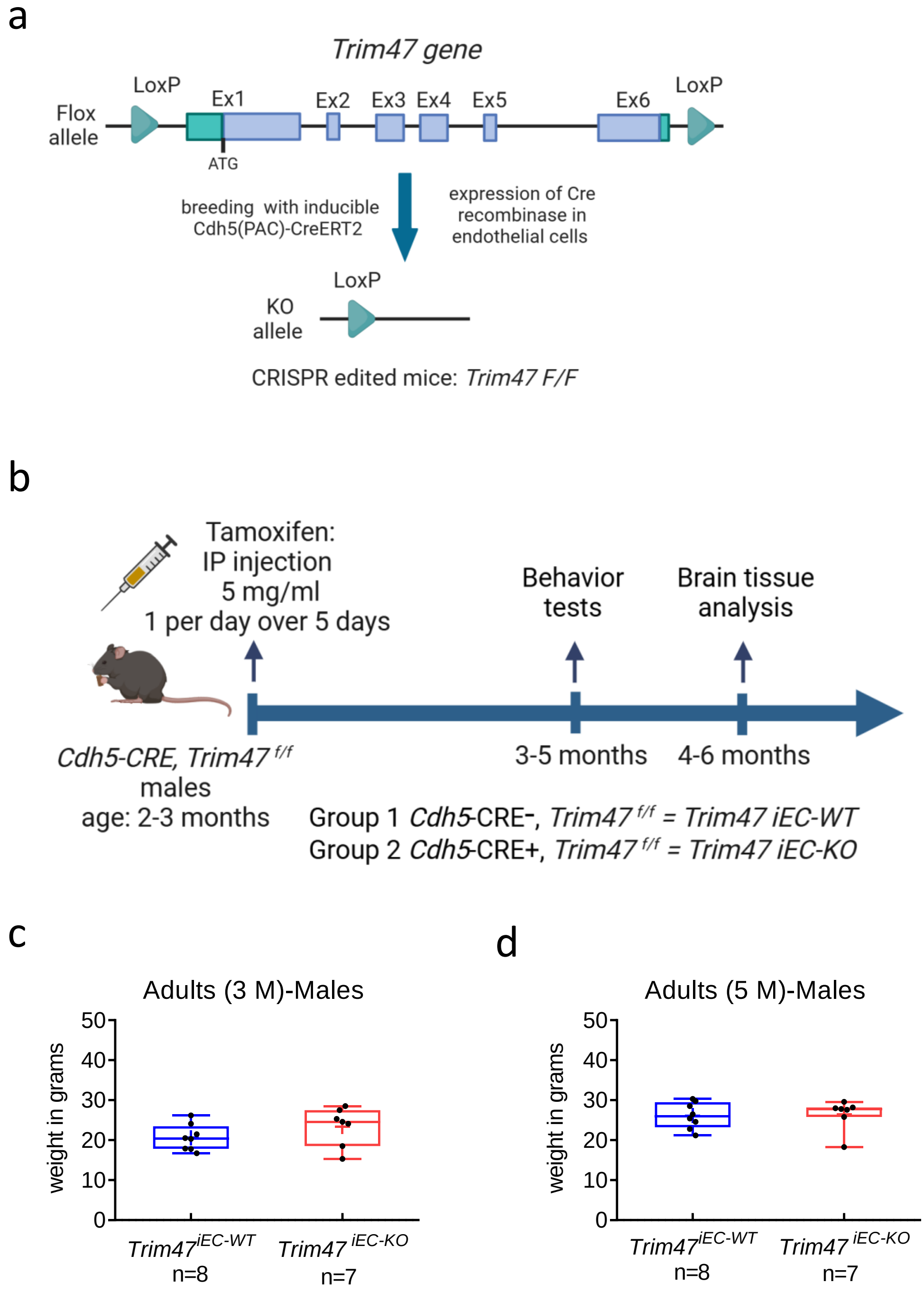


**Supplementary Figure 7**: Generation and characterization of mice deleted for *Trim47* specifically in endothelial cells. **a.** Cartoon depicting the strategy for generating mice deleted for *Trim47* in endothelial cells using *Trim47 F/F* mice (Flox sequences flanking exons 1 to 6) bred with *Cdh5*-Cre mice. **b**. Timeline for induction of *Trim47* deletion in EC following tamoxifen injection in adult mice (2 months). Behavioral tests, brain function assessment and qPCR screening were performed on adults *Trim47* $^{iEC-WT}$ and *Trim47* $^{iEC-KO}$ males. **c.** Body weight expressed in grams of 3 months males (n=8 *Trim47* $^{iEC-WT}$ and n=7 *Trim47* $^{iEC-KO}$). **c.** Body weight expressed in grams of 5 months males (n=8 *Trim47* $^{iEC-WT}$ and n=7 *Trim47* $^{iEC-KO}$).

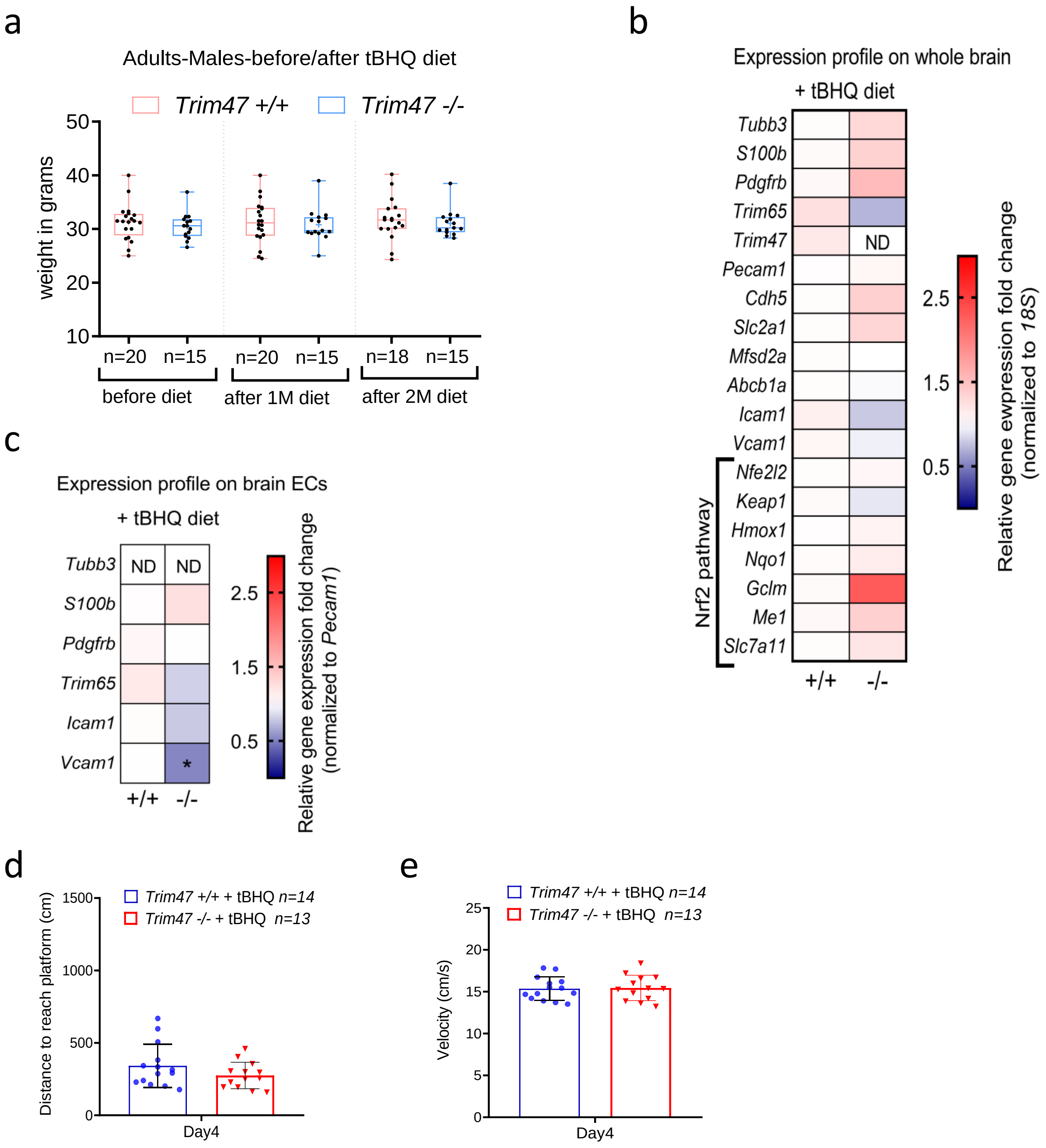


**Supplementary Figure 8**: Generation and characterization of mice deleted for *Trim47* specifically in endothelial cells. **a.** Monitoring of mouse body weight (in grams) before, after 1 month or 2 months of tBHQ diet showing no weight loss over time. **b-c**. qPCR screening on (**b**) whole brain and (**c**) brain EC isolated from *Trim47+/+ and Trim47-/-* males (8-10 months) with tBHQ diet (n=5 replicates/genotype). *: p<0.5; Mann-Whitney. ND: not detectable. **d-e.** Water maze test was performed on *Trim47+/+ and Trim47-/-* males with tBHQ diet (n=14 *Trim47+/+* + tBHQ and n=13 *Trim47+/+* + tBHQ). (**d**) Graph shows the distance to reach platform (expressed in cm) for the day 4. (**e**) Data represent the velocity expressed as cm per second (swimming speed) of each mouse at day 4 during water maze. All Graphical data are mean ± s.d.

| Target | Company | Sequence |
|---|---|---|
| siControl | Eurogentec | SR-CL000-005 |
| siTRIM47 #1 | Eurogentec | CCAGGGACUAUUUCCUCAA55 |
| siTRIM47 #2 | Eurogentec | GCAGCUGUUUGGAACCAAA55 |
| siNRF2 | Eurogentec | AUUGAUGUUUCUGAUCUAUCACU55 |

**Supplementary Table 1**: Sequences of siRNA used for HBMEC and HeLa transfection

| Antibody (host) | Company | Catalogue number | Application and dilution |
|---|---|---|---|
| CD68 (rat) | Biolegend | 137001 | IF (mouse brain, 1/200) |
| Claudin5-488 (mouse) | ThermoFisher | 352588 | IF (mouse brain, 1/200) |
| Fibrinogen (rabbit) | Dako | A0080 | IF (mouse brain, 1/200) |
| GFAP (rabbit) | ThermoFisher | OPA1-06100 | IF (mouse brain, 1/400) |
| Glut1 (rabbit) | ThermoFisher | PA1-1063 | IF (mouse brain, 1/200) |
| HO1 (mouse) | Enzo life Science | ADI-OSA-110 | WB (HBMEC, 1/1000) |
| IBA1 (rabbit) | Fujifilm Wako | 019-19741 | IF (mouse brain, 1/200) |
| Isolectin B4-FITC | Sigma | L 2895 | IF (mouse retina, 1/200) |
| KEAP1 (rabbit) | Abcam | ab227828 | WB (HBMEC, 1/1000), Co-IP (Hek293, 2 µg for 500 µg of protein lysates) |
| Myelin Basic Protein (rat) | Abcam | ab7349 | IF (mouse brain, 1/200) |
| Myc (mouse) | Upstate | 05-419 | WB (Hek293, 1/1000) |
| Myc (mouse) | Millipore | 05-724 | Co-IP (Hek293, 2 µg for 500 µg of protein lysates) |
| NeuN (rabbit) | Millipore | ABN78 | IF (mouse brain, 1/200) |
| NRF2 (rabbit) | Abcam | ab62352 | WB (HBMEC, 1/1000) |
| Podocalyxin (goat) | R&D Systems | AF1556 | IF (mouse brain, 1/400) |
| TRIM47 (rabbit) | Invitrogen | PA5-110521 | WB (HBMEC , 1/1000) |
| TRIM47(rabbit) | Proteintech | 26885-1-AP | WB (mouse brain, 1/1000) |
| α-tubulin (mouse) | Sigma | T5168 | WB (HBMEC, 1/30000) |

**Supplementary Table 2**: List of antibodies used for this study on HBMEC and mouse tissue. IF: Immunofluorescence; WB: Western-Blot, Co-IP: co-immunoprecipitation. Dilution of the antibodies used for each specific application is specified in brackets.

| Target | Forward | Reverse |
|---|---|---|
| *PPIA/cyclophilin* | AGCTAGACTTGAAGGGGAATG | ATTTCTTTTGACTTGCGGGC |
| *TRIM47* | TGAGCAGTCCAAAGTCCTGA | CTACGGCTGCACTCTTGATG |
| *NFE2L2/NRF2* | CACATCCAGTCAGAAACCAGTGG | GGAATGTCTGCGCCAAAGCTG |
| *KEAP1* | CCAACTTCGCTGAGCAGATT | GCTGATGAGGGTCACCAGTT |
| *HMOX1/HO1* | CCAGGCAGAGAATGCTGAGTTC | AAGACTGGGCTCTCCTTGTTGC |
| *NQO1* | GAAGAGCACTGATCGTACTGGC | GGATACTGAAAGTTCGCAGGG |

**Supplementary Table 3**: List of human oligonucleotides used for qPCR

| Target | Forward | Reverse |
|---|---|---|
| *Tubb3/Tuj1* | TCAGCGATGAGCACGGCATA | CACTCTTTCCGCACGACATC |
| *S100b* | CTGGAGAAGGCCATGGTTGC | CTCCAGGAAGTGAGAGAGCT |
| *Gfap* | AACCGCATCACCATTCCTGT | TGGCAGGGCTCCATTTTCAA |
| *Cd11b/Itgam* | TACTTCGGGCAGTCTCTGAGTG | ATGGTTGCCTCCAGTCTCAGCA |
| *Pdgfrb* | GCTAGCTGGTTGGCTAGCTG | CTTCCGGTGTCTAAATGTGGGT |
| *Trim65* | AGGAGCAACGCAGTCGGATTGA | GCCTGCTTCTTGGCTACCTCTA |
| *Trim47* | CTACAGAAACTCGGCTCAGAAGAT | GACTCCGGGTAGTTGATGGG |
| *Pecam1* | TCATTGGAGTGGTCATCGCC | TGTTGGAGTTCAGAAGTGGAGCAG |
| *Cdh5* | CCTGTAGGGAAAGAGTCCATTGTG | ACTTGACCGTGATGTTGGCG |
| *Slc2a1/Glut1* | TCTCTGTCGGCCTCTTTGTT | GCAGAAGGGCAACAGGATAC |
| *Abcb1a* | TCCTCACCAAGCGACTCCGATA | ACTTGAGCAGCATCGTTGGCGA |
| *Mfsd2a* | GCTCTGTCACCTCCTCACTG | ACGTTTCTACATTAGTGTTCCGAG |
| *Nfe2l2/Nrf2* | TAGATGACCATGAGTCGCTTGC | GCCAAACTTGCTCCATGTCC |
| *Keap1* | ATCCAGAGAGGAATGAGTGGCG | TCAACTGGTCCTGCCCATCGTA |
| *Hmox1/Ho1* | CACTCTGGAGATGACACCTGAG | GTGTTCCTCTGTCAGCATCACC |
| *Nqo1* | AGGATGGGAGGTACTCGAATC | AGGCGTCCTTCCTTATATGCTA |
| *Gclm* | AGGAGCTTCGGGACTGTATCC | GGGACATGGTGCATTCCAAAA |
| *Me1* | GTCGTGCATCTCTCACAGAAG | TGAGGGCAGTTGGTTTTATCTTT |
| *Slc7a11* | CTTTGTTGCCCTCTCCTGCTTC | CAGAGGAGTGTGCTTGTGGACA |
| *Icam1* | TGGCCTGGGGGATGCACACT | CCACCGGGCTGTAGGTGGGT |
| *Vcam1* | CGTACACCATCCGCCAGGCA | TAGAGTGCAAGGAGTTCGGGCG |
| *Il1β* | TGGACCTTCCAGGATGAGGACA | GTTCATCTCGGAGCCTGTAGTG |
| *Il6* | CACTTCACAAGTCGGAGGCT | CTGCAAGTGCATCATCGTTGT |
| *Cldn5* | ACGGGAGGAGCGCTTTAC | GTTGGCGAACCAGCAGAG |
| *Cldn11* | GCCTGGAGTGGCCAAGTA | AGATGGTGGCGACAATGG |
| *Ocln* | GTCCGTGAGGCCTTTTGA | GGTGCATAATGATTGGGTTTG |
| *Tjp1* | AAGTTGGCAAGAGAGGAGCC | CAACCGCATTTGGCGTTACA |
| *Plvap* | GTTGACTACGCGACGTGAGATG | AGCTGTTCCTGGCACTGCTTCT |
| *18S* | CGCGGTTCTATTTTGTTGGT | AGTCGGCATCGTTTATGGTC |

**Supplementary Table 4**: List of mouse oligonucleotides used for qPCR

**Supplementary data 5**: Uncropped immunoblots underlying Figures

### Uncropped immunoblots underlying Figure 1d

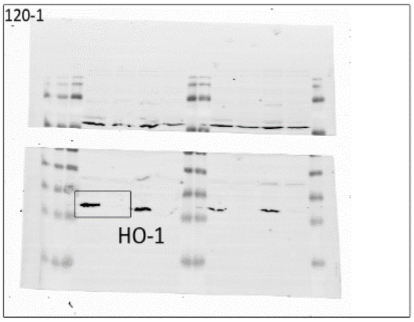


12% Acrylamide Gel

Channel 700nm Low Exposure

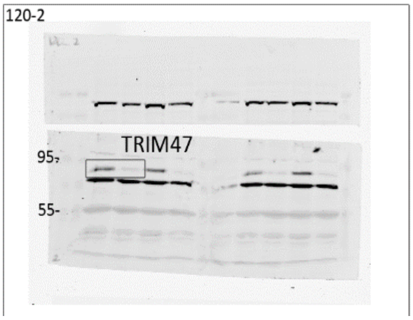


6% Acrylamide Gel

Channel 800nm Low Exposure

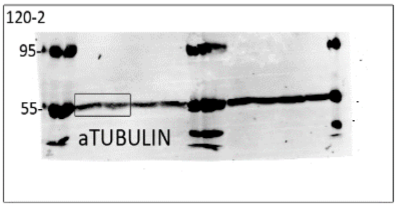


6% Acrylamide Gel

Channel 700nm

### Uncropped immunoblots underlying Figure 2d

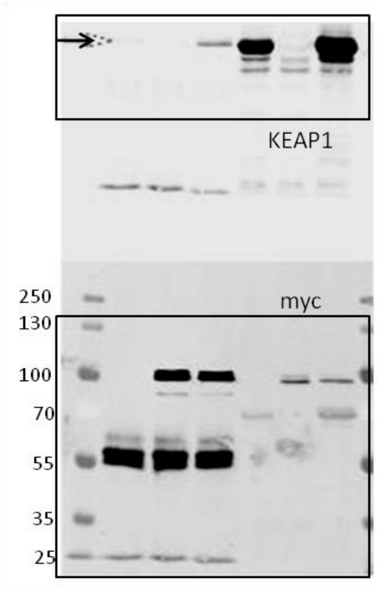


Channel 800nm

Channel 700nm

### Uncropped immunoblots underlying Figure 2e

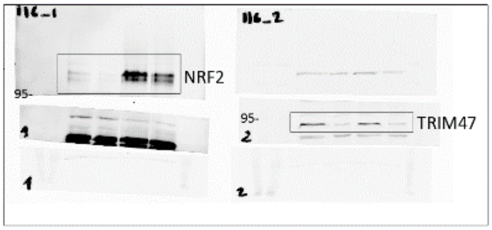


6% Acrylamide

Channel 800nm

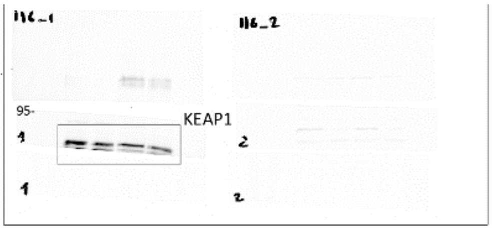


6% Acrylamide

Channel 800nm Low Exposure

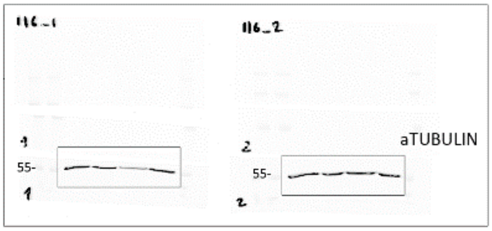


6% Acrylamide

Channel 700nm Low Exposure

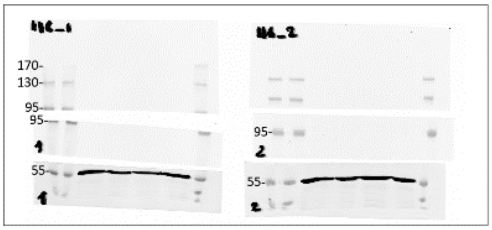


6% Acrylamide

Channel 700nm
